# Decoding the contents of consciousness from prefrontal ensembles

**DOI:** 10.1101/2020.01.28.921841

**Authors:** Vishal Kapoor, Abhilash Dwarakanath, Shervin Safavi, Joachim Werner, Michel Besserve, Theofanis I. Panagiotaropoulos, Nikos K. Logothetis

## Abstract

Multiple theories attribute to the primate prefrontal cortex a critical role in conscious perception. However, opposing views caution that prefrontal activity could reflect other cognitive variables during paradigms investigating consciousness, such as decision-making, monitoring and motor reports. To resolve this ongoing debate, we recorded from prefrontal ensembles of macaque monkeys during a no-report paradigm of binocular rivalry that instigates internally driven transitions in conscious perception. We could decode the contents of consciousness from prefrontal ensemble activity during binocular rivalry with an accuracy similar to when these stimuli were presented without competition. Oculomotor signals, used to infer conscious content, were not the only source of these representations since visual input could be significantly decoded when eye movements were suppressed. Our results suggest that the collective dynamics of prefrontal cortex populations reflect internally generated changes in the content of consciousness during multistable perception.

**One sentence summary:** Neural correlates of conscious perception can be detected and perceptual contents can be reliably decoded from the spiking activity of prefrontal populations.

## INTRODUCTION

One of the most elusive problems in science is to understand the biological basis of consciousness (*1–3*). A seminal paper almost 30 years ago, incited researchers that *“the time is ripe for an attack on the neural basis of consciousness”* and proposed conscious visual perception as a form of consciousness that is within the reach of neuroscience (*4*). Since then, several theoretical treatises (*5–8*) including the frontal lobe hypothesis (*6*), the higher order (HOT) (*8*) and global neuronal workspace (GNW) theories (*5, 9*) have suggested a critical role for the brain’s prefrontal cortex (PFC) in mediating conscious perception. Evidence supporting its involvement comes from functional magnetic resonance imaging (fMRI) (*9, 10*), experience of visual hallucinations upon electrical stimulation of the region (*11, 12*), impaired conscious processing following PFC lesions in patients (*13–18*) as well as intracortical recordings of neural activity (*19–22*). In contrast, alternative theories like the IIT (integrated information theory) identifies the neural elements mediating consciousness as the system having maximal internal cause-effect power (*23*). Its proponents among others have suggested that the neural substrates underlying conscious perception can be traced to a “posterior cortical hot zone”, with PFC being primarily critical for processing the behavioral and cognitive consequences of conscious perception (*24–26*) like task demands and monitoring, introspection and motor reports, rather than consciousness per se (*26–29*). For example, frontal cortex was found to be dramatically more active during motor reports, when blood-oxygen-level-dependent (BOLD) fMRI signal modulation was compared between reported and unreported spontaneous changes in the content of consciousness (*27, 30, 31*). Additionally, reduced frontal activation accompanied inconspicuous and unreportable switches in perception, when contrasted against a condition, where perceptual changes were easily discernible (*29*). Together, these reports could suggest that frontal activity is related to consequences of perception, thus casting doubt on its role in conscious content representations (*13, 25, 26, 32*). However, the univariate fMRI analysis comparing report vs no-report conditions (*27*) as well as the indirect nature and limited spatial resolution of BOLD fMRI signal compared to neuronal recordings leaves open the possibility that prefrontal ensembles do reflect the content of consciousness even without report requirements.

We examined this hypothesis by simultaneously probing the discharge activity of large neural populations in the inferior convexity of the macaque PFC with multielectrode arrays during a no-report binocular rivalry (BR) paradigm. BR belongs to the family of multistable perceptual phenomena (*7, 33, 34*), which allow a dissociation of conscious perception from sensory input, by inducing in an observer spontaneous fluctuations in the content of consciousness without a change in sensory stimulation. BR instigates these perceptual fluctuations through presentation of incongruent, dichoptic visual input to corresponding retinal locations, resulting in stochastic, endogenously driven alternations in subjective perception. For a certain duration, only one of the two images is consciously experienced, while the other is perceptually suppressed (*33, 34*). Similar to other paradigms utilized in investigating conscious perception, the standard practice in BR requires humans and macaques to manually report their percepts (e.g. by pressing levers). Therefore neural activity related to consciousness could be conflated with signals related to its consequences such as voluntary motor reports, decision making and introspection (*35–37*). Objective indicators of perception provide a solution to this problem. For example during rivalry between opposing directions of motion, the polarity of optokinetic nystagmus (OKN) reflex, a combination of smooth pursuit and fast saccadic eye movements, is known to be tightly coupled to the reported direction of motion (*38–41*). Thus, the reflexive nature of OKN can be exploited as an objective measure of changes in the content of consciousness (*41*), removing any confounds in the neural activity originating from voluntary motor reports.

We therefore combined neural ensemble recordings with no-report BR between opposing directions of motion and found that prefrontal neural activity reflects the OKN-inferred content of visual consciousness during spontaneous perceptual switches. Monitoring simultaneously large neural populations allowed us to observe for each spontaneous perceptual transition, the collective dynamics of neuronal ensembles representing two percepts that compete for access to consciousness. Conscious content could be successfully decoded from feature specific ensemble vectors in single instances of internally generated perceptual transitions, thus reinforcing the role of prefrontal populations in mediating conscious perception.

## RESULTS

We exposed two rhesus macaques to a no report BR paradigm, which consisted of two trial types, physical alternation (PA) and binocular rivalry (BR) (see Figure 1A and methods). Each trial started with a fixation spot cueing the animal to initiate fixation, which lasted ∼300 milliseconds, followed by an upward or downward drifting stimulus, presented monocularly for 1 or 2 seconds. After this initial phase, during BR trials, a second stimulus drifting in the opposite direction was presented to the contralateral eye, typically inducing perceptual suppression of the first stimulus, a phenomenon known as binocular flash suppression (BFS) (*42, 43*). Following this period, visual competition ensued, resulting in spontaneous perceptual switches between the two opposing directions of motion. In contrast, PA trials consisted of exogenously driven changes in perception by alternating monocular presentations of the same stimuli used for instigating BR.

**FIGURE 1.**
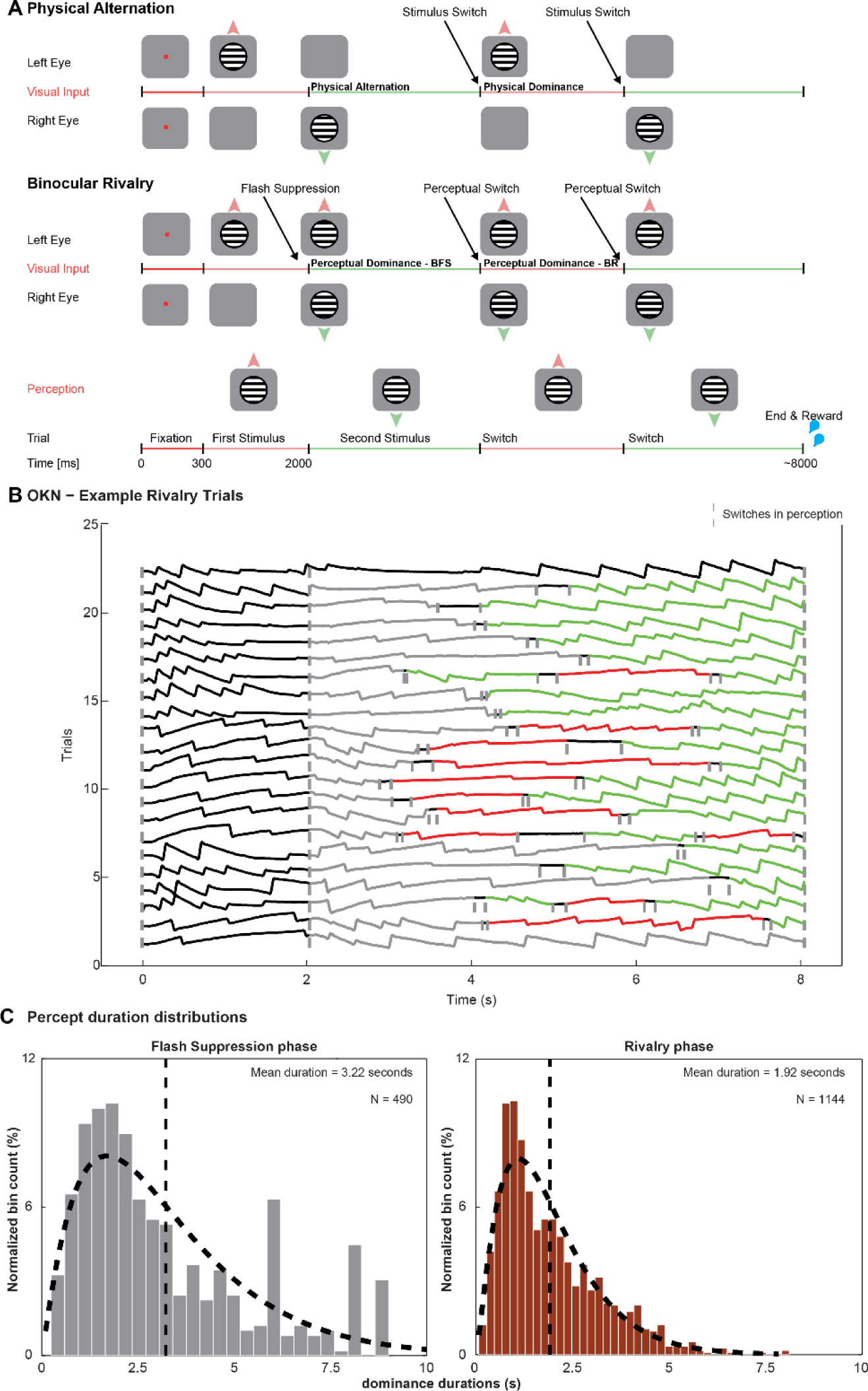
Binocular rivalry paradigm and behavior (A). The task consisted of two trial types, namely, the physical alternation (PA) trials and binocular rivalry (BR) trials. Both trial types started with the presentation of a fixation spot, cueing the animal to initiate fixation. Upon successful fixation for 300 milliseconds, a drifting sinusoidal grating was monocularly presented. After 1 or 2 seconds, the first stimulus was removed and a second grating drifting in the opposite direction was presented in the contralateral eye during PA trials. During BR trials, the second stimulus was added to the contralateral eye without removing the first stimulus, inducing perceptual suppression of the first stimulus (Flash Suppression). After this period, visual input alternated between upward and downward moving gratings during PA trials (Stimulus Switch). During BR trials, the percept of the animal could randomly switch between the discordant visual stimuli (Perceptual Switch). Note that perceived direction displayed in the bottom row schematic is identical, even though the underlying visual input is monocular in PA and dichoptic during BR. (B) OKN elicited during example BR trials from one recording session. The gray vertical dashed line denotes the beginning of the flash suppression phase. Subsequent dominance phases are color coded and their beginning and end are marked with shorter grey dashed lines. Note that on the last example trial the flash suppression resulted in a prolonged continuous suppression of the previously presented direction of motion, while on the first trial, flash suppression was not effective and the initially presented direction of motion remained dominant. (C) Perceptual dominance distributions during flash suppression and rivalry phases could be approximated well with a gamma distribution.

The rivaling, oppositely drifting stimuli elicited optokinetic nystagmus (OKN), (Figure 1B), which served as a reliable indicator for the contents of perception (*38–41*). BFS resulted in a switch in the polarity of the OKN, indicating perceptual dominance of the newly presented direction of motion (marked in grey). Following this initial period of externally induced perceptual suppression, OKN polarity could be observed switching again, occasionally more than once until the end of the trial, indicating spontaneous, internally driven, changes in conscious contents. We evaluated the onset and offset of perceptual dominance periods in every trial based on the stability of the OKN pattern (see methods). These well-defined epochs of stable perceptual dominance during both BFS and BR displayed a gamma distribution (Figure 1C), typical of multistable perception dynamics (*44*) with an average dominance duration of 3.22 ± 0.102 and 1.92 ± 0.037 seconds respectively.

We targeted the inferior convexity of the PFC (Figure 2A and methods), where neurons display selective responses to complex visual stimuli as well as direction of motion (*45–47*). Neural representations of conscious content are directly related to such feature selectivity. If neurons during BR reliably increase their firing rate each time their preferred stimulus is perceived and suppress their activity, when it is perceptually suppressed, then they explicitly represent conscious content. Figure 2B displays discharges of four such, simultaneously recorded prefrontal units and OKN during a single BR trial. Two units (33 and 119) fired more when downward drifting stimulus was presented while the other two (44 and 167) displayed stronger modulation for the opposite direction of motion during a PA trial (Supplementary Figure 2.1). Spiking activity of these units was also correlated with subjective changes in conscious perception in a BR trial, for both externally induced perceptual suppression (BFS) and a subsequent spontaneous switch in conscious content (BR) (Figure 2B and D).

**FIGURE 2.**
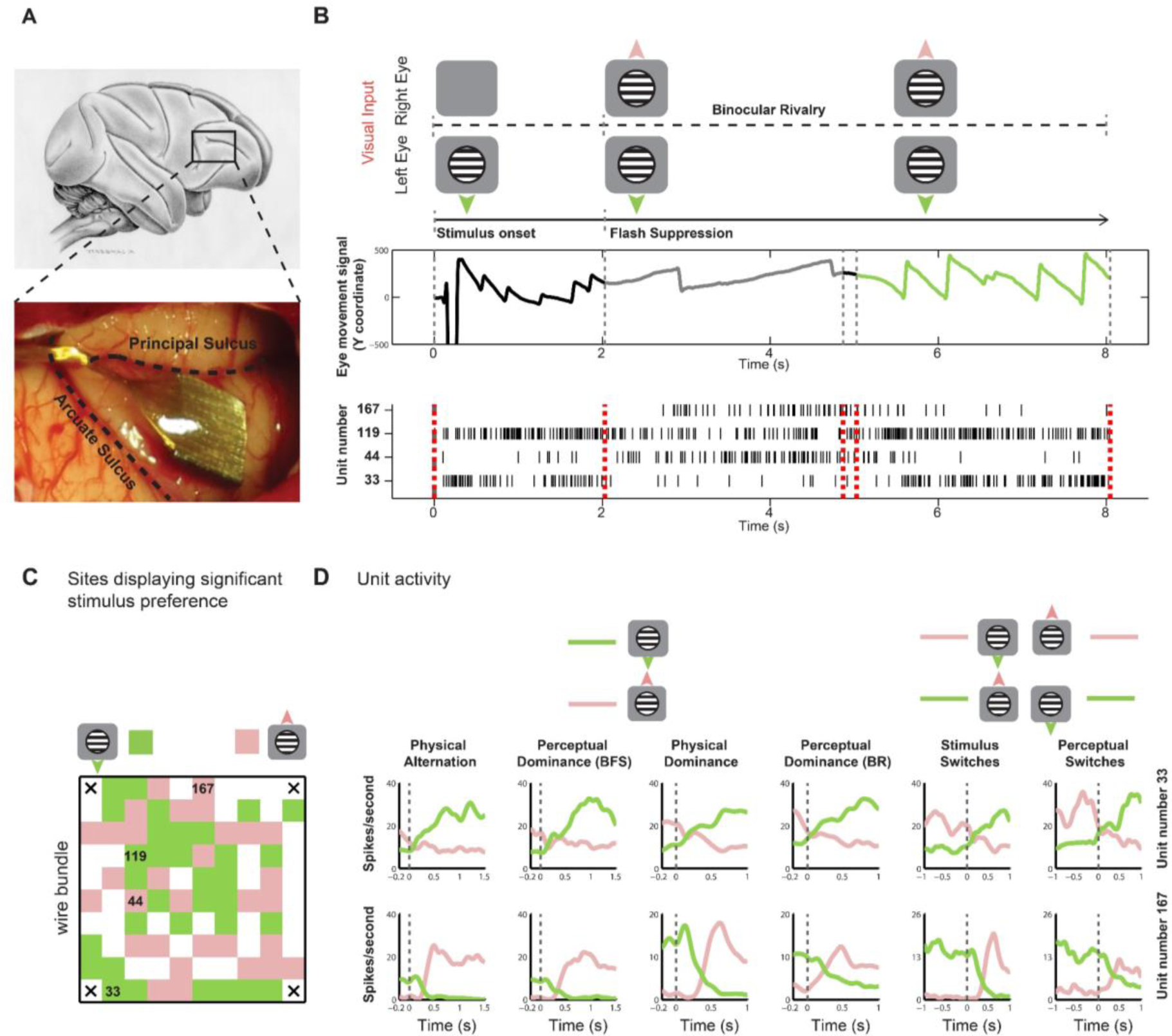
Example unit responses (A) The location of the implanted Utah array in the inferior convexity of the PFC on a schematic macaque brain and in one of the animals. (B) Visual input, OKN and corresponding spiking activity during an example BR trial. The trial started with monocular presentation of downward drifting grating. An upward drifting grating was added to the contralateral eye 2000 ms later, which resulted in perceptual suppression (flash suppression) of downward motion, inferred from a change in the OKN direction. Externally induced perceptual suppression lasted for ∼3000 ms after which a spontaneous switch reinstated the perception of the downward motion. In the spike raster plot, while unit numbers 33 and 119 display strong spiking activity when downward drifting grating is perceptually dominant, unit numbers 44 and 167 respond stronger when upward drifting grating is perceived. (C) Projection of all sites with significant stimulus preference during the flash suppression phase of the PA trials on the array for one recording session. The location of the units presented in (B) are marked. Green and pink pixels reflect sites, where spiking activity (unsorted spikes recorded from a given electrode), responded more to upward or downward drifting gratings respectively. (D) Average spike density functions of two units recorded (unit 33, selective for downward motion in the upper row, unit 167, selective for upward motion in the lower row) simultaneously in the PFC during PA and BR trials. Pink and green colors in the first four columns correspond to the response elicited by presentation or perception of downward and upward drifting grating respectively. In the last two columns, we plot the activity elicited during a stimulus or perceptual switch from downward to an upward drifting grating (pink) and vice versa (green). The activity of both units is very similar during PA and BR, thus displaying robust perceptual modulation.

We analyzed separately the spiking activity during perceptual dominance and suppression periods either (i) externally induced during BFS, or (ii) brought about by an endogenous spontaneous switch, in BR. Selectivity of neural activity was analyzed both before and after such perceptual switches and compared to selectivity in corresponding temporal phases from PA trials (see methods). Figure 2D displays the average spike density functions of units 33 (preferring downward motion) and 167 (preferring upward motion). The two units were recorded simultaneously from distant electrodes on the array (Figure 2C) and displayed robust modulation during both PA and BR trials with spiking activity switching reliably for both externally induced (Wilcoxon rank sum test, during PA trials (temporally analogous phase to flash suppression dominance) for unit 33, p_PA-33_ = 2.82*10^-14^ and for binocular flash suppression phase during BR trials, p_BFS-33_ = 2.39*10^-6^; for unit 167, p_PA-167_ = 1.13*10^-17^ and p_BFS-167_ = 1.20*10^-9^) and internally driven perceptual switches (Wilcoxon rank sum test, during PA trials (temporally equivalent phase to rivalry dominance) for unit 33, p_PA-33_ = 8.72*10^-15^ and perceptual dominance during BR trials, p_BFS-33_ = 7.18*10^-8^; for unit 167, p_PA-167_ = 1.49*10^-4^ and p_BFS-167_ = 7.8*10^-3^). The recorded sites displaying similar stimulus preference formed clusters throughout the 16mm^2^ recorded area of the inferior prefrontal convexity (Figure 2C and Supplementary Figure 2.1).

We compared the stimulus selectivity of all recorded units (n = 987 and see methods) during subjective perception in BFS and BR with their selectivity during purely sensory, monocular stimulus presentations in PA trials, using a d’ index (see methods) (*20, 48*). A large majority of units exhibiting significant stimulus selectivity in PA (see Methods), fired more when their preferred stimulus was perceived compared to its perceptual suppression during BR trials (BFS - 85.38 % (292/342) and BR - 76.09 % (277/364)), with 53.8 % (184/342) and 40.38% (147/364) of them being significantly modulated, respectively (Wilcoxon rank sum test, p<0.05, also see Table 1) (Figure 3A), suggesting that ongoing perceptual content is robustly encoded in PFC. Moreover, compared to earlier visual regions, where many units display significant modulation during perceptual suppression of their preferred stimulus, which were proposed as part of an inhibitory mechanism independent of the mechanisms of perception, suggestive of non conscious processing (*49*); such units were a small minority in PFC (BFS - 2.92 % (10/342) and BR - 4.12 % (15/364)). Furthermore, many units displayed significant preference only during BFS (26.51 %, 70/264) and BR (34.4 %, 85/247), suggesting that individual prefrontal units contribute more reliably to conscious perception during visual competition.

**FIGURE 3.**
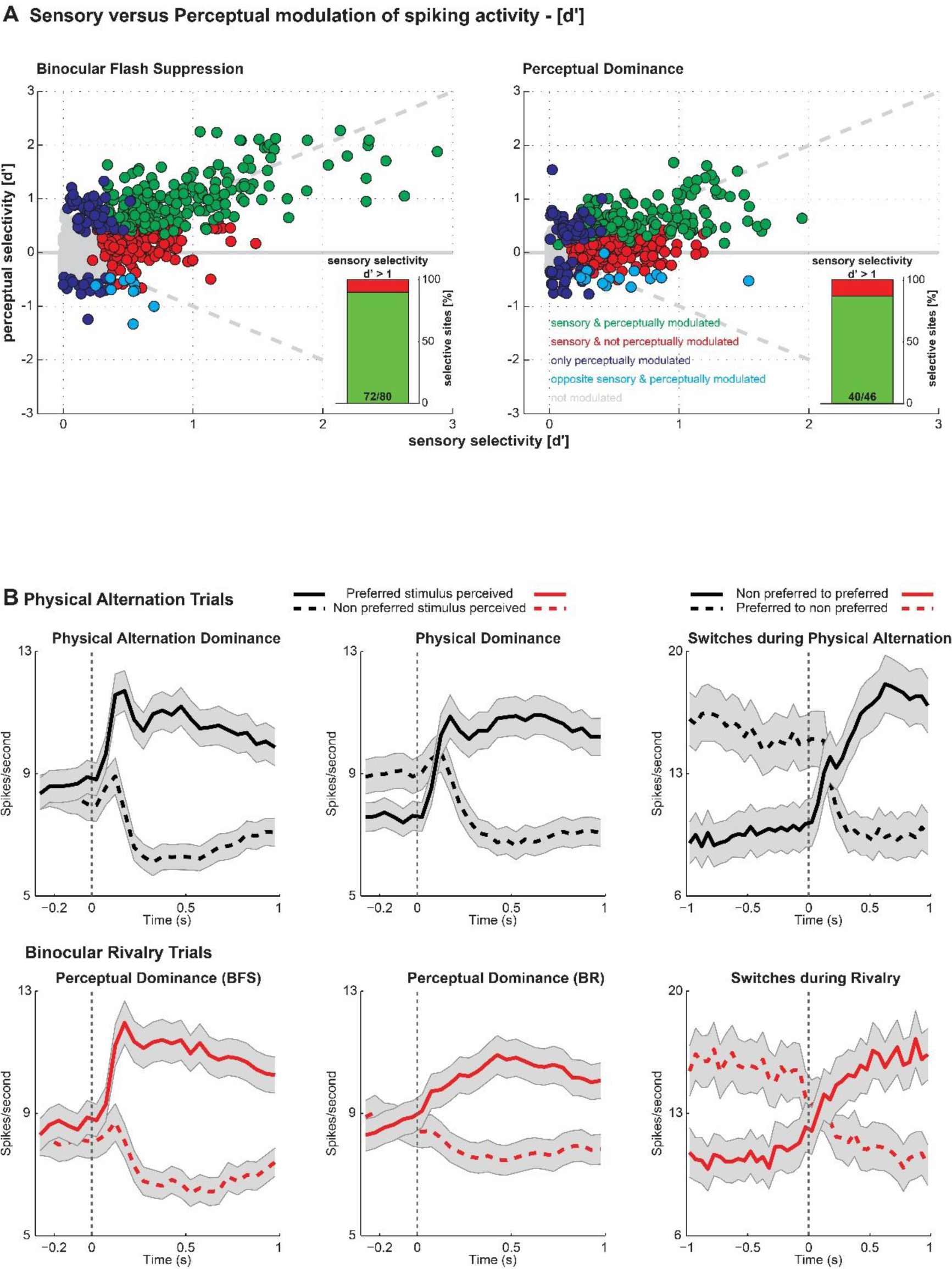
Sensory (PA) versus perceptual (BR) modulation of spiking activity. (A) Scatter plot of sensory vs. perceptual selectivity (d′) for all units (dots) across all datasets for BFS and BR. Units showing no significant modulation in PA or BR trials are displayed in grey, those with significant modulation during both conditions in green, units which display significant preference only during PA trials in red and units displaying significant modulation only during BR trials are displayed in blue. In cyan the small percentage of units which fired more when their preferred stimulus was perceptually suppressed across the two conditions. The proportion of perceptually modulated units for both BFS (90%) and BR (86%) increased as a function of sensory selectivity strength (insets showing perceptual modulation for d’>1). (B). Average population spiking activity in PA and BR. The population activity averaged across all units which were significantly modulated during PA (upper row) or BR (lower row) trials and preferred the same stimulus is plotted for the flash suppression (left), the perceptual dominance (middle) phase and switches (right) during BR and temporally matched phases in PA. Displayed are two traces of population activity, one, calculated when the unit’s preferred stimulus was dominant (thick lines) and the second, when the unit’s preferred stimulus was suppressed because of the dominance of its non-preferred stimulus (dashed lines). Population activity reliably followed phenomenal perception during perceptual transitions brought about exogenously with flash suppression as well as endogenously driven during BR. A remarkable similarity in population activity across the two trial types indicates strong and robust perceptual modulation.

**Table 1.** Number (N) and proportion (%) of significantly modulated units during the BR paradigm

| **Trial Type** | **Physical Alternation** |  | **Binocular Rivalry** |  |
| --- | --- | --- | --- | --- |
| **Temporal Phase** | **Physical Alternation** | **Sensory Dominance** | **BFS Dominance** | **BR dominance** |
| **N of significantly modulated units** | **342** | **364** | **264** | **247** |
| **N and % of significantly modulated units\* in PA, with similar average stimulus preference in BR** | **292/342**<br>**85.38%** | **277/364**<br>**76.09%** |  |  |
| **N and % of significantly modulated units\* in BR, with similar stimulus preference in PA** |  |  | **229/264**<br>**86.74%** | **199/247**<br>**80.56%** |
| **N of significantly modulated units in PA with a $d' > 1$** | **80** | **46** |  |  |
| **N and % of significantly modulated units\* in PA ($d' > 1$), with similar preference and significantly modulated in BR (ALL)** | **72/80**<br>**90%** | **40/46**<br>**86.96%** |  |  |

\*significantly modulated here in a particular condition refers to units displaying significantly stronger activity to one of the two stimuli using a Wilcoxon rank sum test at an alpha value of 0.05.

Over all, the recorded units displayed considerable heterogeneity in stimulus preference strength (d’) during BFS (PA - 0.3985 ± 0.0131, BFS - 0.4471 ± 0.0133) and BR (PA - 0.3075 ± 0.0098, BFS - 0.2719 ± 0.0082) (Figure 3A). Importantly, selectivity strength was a critical factor determining significant perceptual modulation in BFS and BR. For units with strong sensory selectivity (d’ > 1), around 90% were significantly perceptually modulated (Wilcoxon rank sum test, p < 0.05) (90 % (72/80) for BFS and 86.96 % (40/46) for BR). This indicates that the percentage of perceptually modulated units in the PFC is remarkably similar to the temporal lobe (*50*), thus suggesting that there are at least two cortical regions, where neuronal activity explicitly represents conscious contents. Furthermore, these results show that the activity of prefrontal units correlates with internally driven switches in the subjective perception of more simple visual features like direction of motion, in addition to the externally induced perceptual suppression of faces and more complex stimuli (*20*).

Plotting the population spiking activity averaged across all units which displayed significant modulation and similar preference across the first monocular switch phase of PA and the temporally corresponding flash suppression phase of the BR trials, revealed a strong, early peak response followed by a long sustained response when a preferred stimulus was presented and a dramatic suppression in activity during presentation of the non-preferred stimulus (Figure 3B, upper row). The average population activity during the BFS phase displayed robust perceptual modulation firing more when a preferred stimulus was perceived, and less when a preferred stimulus was suppressed by a non-preferred stimulus stimulating the contralateral eye (Figure 3B, lower row). Similarly, reliable perceptual modulation was observed, when stimuli were perceived following spontaneous changes in perception during BR (Figure 3B middle column) as well as around spontaneous perceptual switches (Figure 3B, last column, also see Supplementary Figures 3.1 and 3.2). Similar results were obtained when neural activity in PA was aligned to OKN changes (see methods), as in BR trials (Supplementary Figures 3.3, 3.4, 3.5, and Table 2).

**Table 2.** Number and proportion of significantly modulated units during the BR paradigm (Physical alternation trials aligned to the change in the OKN direction)

**(Physical alternation trials aligned to the change in the OKN direction)**
| **Trial Type** | **Physical Alternation** | **Binocular Rivalry** |
| --- | --- | --- |
| **Temporal Phase** | **Sensory Dominance** | **BR dominance** |
| **N of significantly modulated units** | **408** | **247** |
| **N and % of significantly modulated units\* in PA, with similar average stimulus preference in BR** | **324/408**<br>**79.41%** |  |
| **N and % of significantly modulated units\*in BR, with similar stimulus preference in PA** |  | **199/247**<br>**80.56%** |
| **N of significantly modulated units in PA with a $d' > 1$** | **68** |  |
| **N and % of significantly modulated units\* in PA ($d' > 1$), with similar preference and significantly modulated in BR (ALL)** | **49/68**<br>**72.05%** |  |

Probing the PFC with multi-electrode arrays allowed us to monitor simultaneously, feature specific ensembles, displaying preferential responses to stimuli drifting in opposite directions. We therefore examined the population code for single instances of different types (i.e., upward to downward and downward to upward motion) of exogenous stimulus and endogenous, spontaneous perceptual transitions. Prefrontal ensemble activity correlated with both exogenous stimulus changes in PA and subjective changes in perceptual content during BR trials (Figure 4A). We utilized a multivariate decoding approach (*51*) to assess the reliability with which we could predict conscious perceptual contents from ensemble activity on single cases of perceptual transitions (see methods). During PA switches, the classifier discriminated between the two stimuli strongly above chance (50%), and generalized across the total duration of a given stimulus presentation (Figure 4B, upper row) suggesting a static population code (*52*). Similarly, a classifier trained on BR activity also discriminated between periods of perceptual dominance and suppression for the two competing stimuli and generalized around perceptual switches (Figure 4B, lower row), similar to that observed during stimulus switches. Importantly, strong temporal generalization of the classifier trained and tested across PA and BR before and after a switch, indicates an invariance in the population code representing sensory input and its subjective experience (*51*). This cross trial type generalization was highly significant (permutation test, p<0.002, see methods) when it was carried out during two temporal windows (400 ms), before (-200 ms to - 600) and after (200ms to 600 ms) a switch (Figure 4C). Similarly strong decoding of perceptual content was possible in individual datasets (Supplementary Figure 4.1). Together, these results suggest that the prefrontal population code underlying sensory input and subjective perception is not only similar, but also reliable and robust. Similar results were obtained when PA trials were aligned to the OKN change instead of the digital pulses for stimulus presentation, (Supplementary Figure 4.2).

**FIGURE 4.**
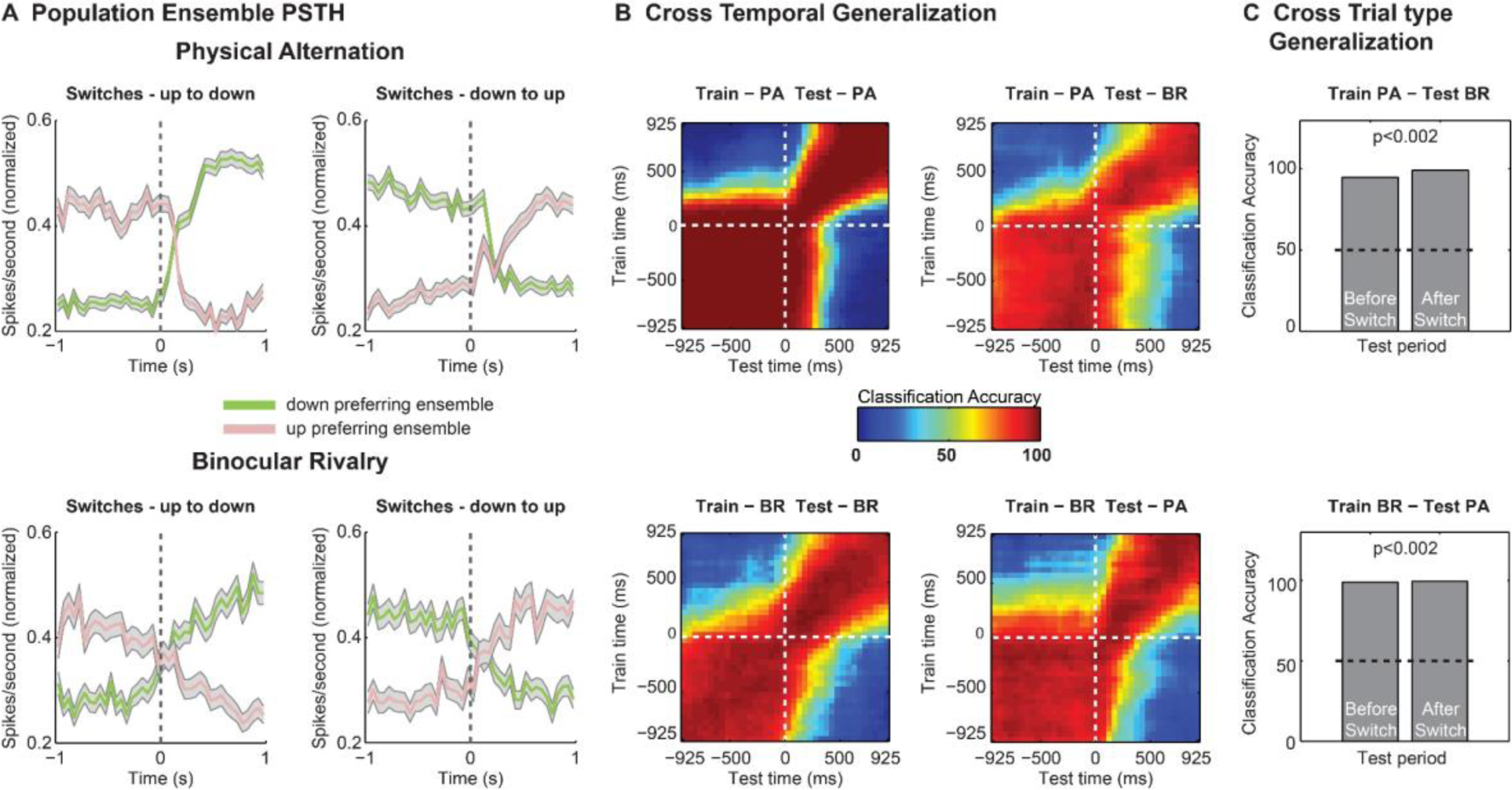
Decoding the contents of conscious perception from simultaneously recorded prefrontal ensembles. (A) Normalized spiking activity of down (green) and up (pink) preferring ensembles of units during up to down or down to up, PA (upper row) and BR (lower row) switches showing reliable modulation of neuronal ensembles during both external stimulus changes and internally generated switches in conscious perception. (B) Cross-temporal decoding of stimulus contents around switches in perception during PA and BR trials and generalization across the two. Classification accuracy was computed for each pair of train and test time windows around a switch (see methods) in steps of 50 ms, using 150 ms bins. (C) Cross trial type generalization was highly significant (permutation test, p < 0.002), suggesting that the underlying population code is invariant to the trial type, and therefore encodes perceptual contents.

Finally, given that in our experiments the OKN is tightly linked to perceptual content, we dissociated neural activity related to oculomotor processes from activity related to visual input. For a majority of the recorded units (n = 747), we estimated their preference to direction of motion in a control experiment during two paradigms, namely fixation Off and fixation On. During fixation Off, the presentation of visual motion elicited OKN similar to BR, while during fixation On, the eye movements were suppressed since macaques were required to fixate a centrally presented spot (Supplementary Figure 5.1 and Supplementary Figure 5.3). We focused our analysis on the upward and downward motion directions, used for instigating rivalry, to make a direct comparison with BR. We found that a majority of the units displaying significant stimulus selectivity across the two control paradigms retained their stimulus preference (fix On - 69.56 %, 48/69; fix Off - 56.25 %, 81/144). Only a small percentage of units (fix On - 14.49 %, 10/69 fix Off - 6.94 %, 10/144; Wilcoxon rank sum test, p <0.05) exhibited a significant preference to stimuli with opposing motion content across the two paradigms (Supplementary figure 5.2 for typical tuning curves). Ensemble population PSTHs (see methods) of significantly modulated units during fixation Off or fixation On preferring the same motion direction in both paradigms are displayed in Figure 5A. In both paradigms, average firing rate increased when a preferred motion direction was presented and decreased in response to the non preferred visual input. We investigated if a classifier trained on neural responses to stimuli which elicited OKN could reliably predict the stimuli, when they were viewed with the eye movements suppressed, and vice versa. We observed strongly above chance (50%) decoding accuracy of the classifier during both conditions (Figure 5B). Importantly, a classifier trained on individual paradigms could generalize across them (Figure 5C) and decode with significant accuracy (permutation test, p<0.002, see methods) thus suggesting that prefrontal ensemble activity contains stimulus information, and is not just driven by the eye movements. Similar results were obtained when decoding analysis was performed using the trials from the fixation On paradigm, where any eye movements within the fixation window were further controlled (see methods)(Supplementary Figure 5.4). These findings are in line with previous work suggesting that frontal cortex responds to visual motion both in the presence and absence of OKN (*47, 53*) and suggest that motion content signals contribute to the activity of the tested population. Neurons in this prefrontal region reflect a mixture of perceptual and oculomotor signals (*54*) and are selective for motion stimuli even when the monkeys fixate (*47*). Such comodulation was recently reported in V4 and inferotemporal cortex where microsaccades were found to contribute in attention related neuronal responses (*55*). Future investigations could ascertain, if a similar mechanism is relevant for prefrontal responses in BR.

**FIGURE 5.**
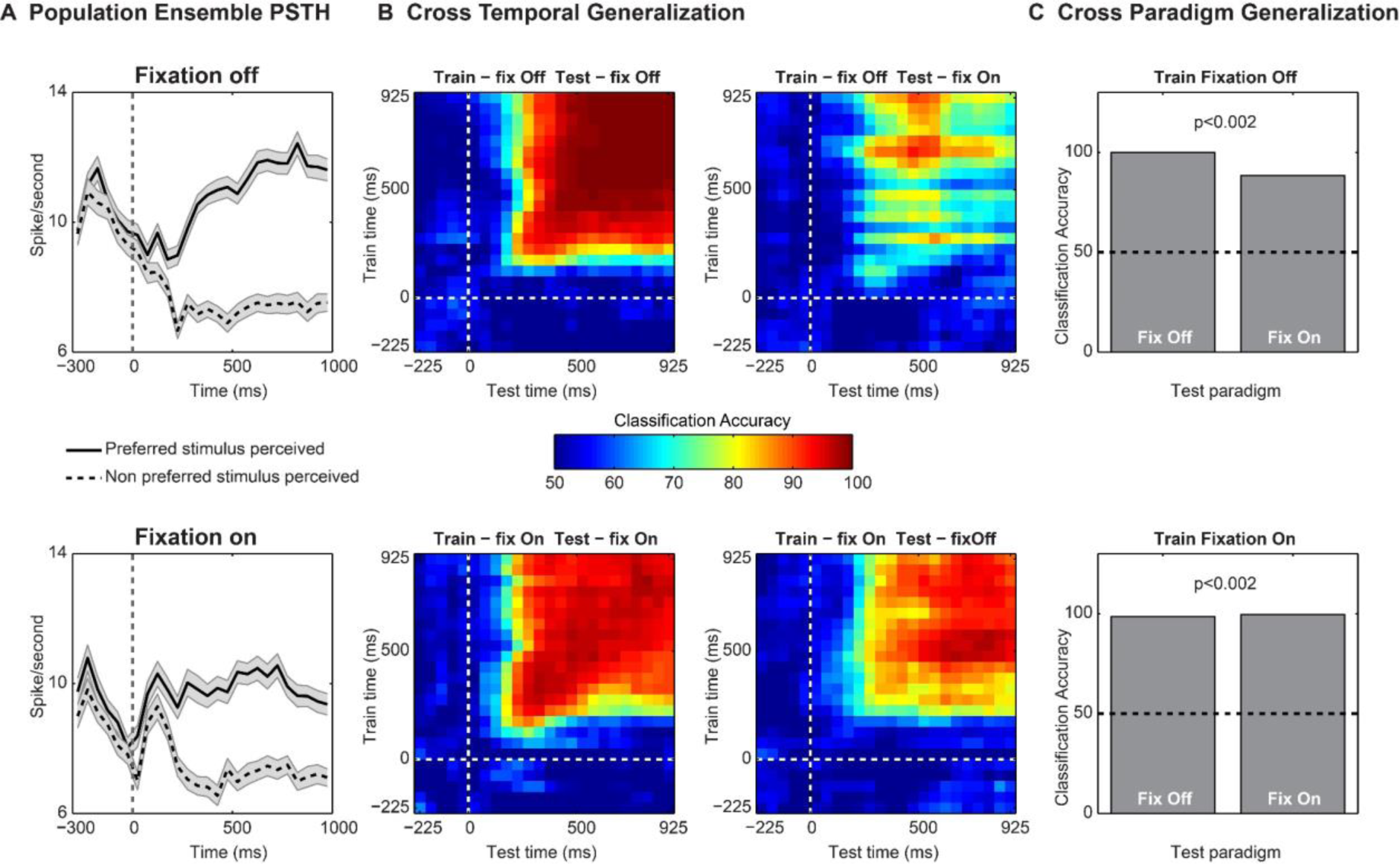
Invariance of the population code to motion content in the presence and during the suppression of OKN eye movements assessed with multivariate pattern analysis. (A) Ensemble population spiking activity (see methods) during the fixation Off and fixation On paradigm for units which were significantly modulated in either of the two paradigms, and preferred the same motion direction (B) Cross-temporal decoding of stimulus contents during the two paradigms. Decoding accuracy was tested for each pair of train and test time windows during the two paradigms as well as across them, with binning parameters similar to figure 4 (C) The cross paradigm invariance of the population code was tested by training a classifier on activity in one paradigm and testing on the other, for a single bin of 400 ms (starting 400 ms post stimulus onset) during the presentation of the visual stimulus. We observed significant (permutation test, p < 0.05) cross-task generalization accuracy, thus suggesting that the underlying code is largely invariant to the presence of large OKN, and encodes stimulus motion contents.

## DISCUSSION

These results suggest that feature selective units in the primate PFC reliably reflect the dynamics of internally generated changes in the content of subjective perception even without voluntary perceptual reports. While addressing an ongoing debate between GNW and IIT about the neural correlates of conscious perception in the PFC (*13, 25, 26, 32, 56*), we demonstrate that the contents of subjective experience can be reliably decoded from the activity of prefrontal ensembles during single instances of internally driven transitions in conscious perception.

BR offers a distinct advantage over other paradigms of visual consciousness such as BFS or visual masking due to the stochastic, internally driven changes in the subjective perceptual content in the absence of any concomitant changes in visual stimulation (*34, 49, 57*). Hence, it confers a unique opportunity to observe neural dynamics contemporaneous with spontaneous changes in the contents of subjective experience. When paired with electrophysiological investigations of the non human primate visual system (*57, 58*), BR has revealed that the proportion of feature selective neurons reliably reflecting conscious content increases as one progresses in the visual cortical hierarchy from early visual areas (*48, 59, 60*) to later temporal regions (*20, 21, 50*). Recent single unit recordings in human medial frontal and anterior cingulate cortex areas during BR found non-selective modulation of neural activity before spontaneous perceptual transitions, suggesting that some frontal cortical areas might reflect the prerequisites of conscious perception than conscious content per se (*22*). In contrast, our results demonstrate conscious content representations in a subregion of the macaque lateral PFC, where cells are selective for faces, complex visual objects and direction of motion (*45*– *47, 61, 62*) and reciprocally connected with the inferotemporal cortex (*63*). Importantly, previous electrophysiological studies in the PFC during conscious perception either utilized a motor report (*19, 21, 22, 64*), and were therefore conflated by consequences of conscious perception, or investigated perceptual modulation among neurons selective to faces and complex objects with a no-report BFS paradigm. In BFS, perceptual dominance and suppression are externally induced due to an abrupt and strong change in the feedforward input and not endogenously driven as in BR, wherein neural activity modulations could contribute causally towards changes in conscious perception (*20*). Hence, our results collected during unreported spontaneous transitions in conscious perception unequivocally demonstrate the existence of prefrontal representations of conscious content.

Our findings are in sharp contrast to the conclusions of recent imaging studies showing reduced involvement of the PFC in conscious perception (*27–29*). However, constraints in the spatiotemporal resolution of the BOLD signal and its complex relationship with neural activity limit the interpretations from imaging data, especially so, when null findings are reported (*65, 66*). Such limits in spatiotemporal resolution are particularly relevant to the frontal cortex, where individual neurons often display a high degree of mixed selectivity (*67, 68*) or distinct temporal patterns of activity during perceptual paradigms (*69*). For example, we find that units displaying preferential responses to opposite directions of motion are frequently distributed in close proximity (∼0.4mm) throughout the electrode array (Figure 2C and supplementary figure 2.3). Such spatial variability of stimulus selectivity remains difficult to capture with fMRI. A recent attempt with high-field fMRI offering better spatial resolution could identify clusters activated by competing perceptual representations and reported a relatively low correlation between sensory and perceptual representations in early visual areas, confirming earlier electrophysiological studies (*70*). In contrast, recent work utilizing fMRI in conjunction with multivariate pattern analysis revealed neural correlates of consciously perceived location in anterior brain regions such as the frontal cortex, beyond early visual areas (*71*). Such approaches hold great promise in providing whole brain representations of conscious content.

Utilizing multivariate pattern analysis for decoding the contents of conscious perception from neuronal ensembles in the PFC lays the foundation for a comparative approach using direct neuronal recordings, aimed at investigating the population code subserving conscious contents across cortical regions. It may further help elucidate the details of the mechanism responsible for inducing spontaneous changes in the content of consciousness. In summary, our results demonstrate that prefrontal ensemble activity explicitly reflects internally generated changes in conscious contents since only a miniscule percentage of units fired significantly more when their preferred stimulus is perceptually suppressed. They, therefore lend support to theoretical approaches such as the GNW and HOT, which attribute an essential role for the PFC in mediating consciousness in general and conscious perception in particular (*8, 72, 73*). Interestingly, apart from conscious contents, PFC was recently shown to control also the level of consciousness in rodents (*74*), suggesting a more general role of this area in awareness. While GNW and HOT have recently received criticism because of this region’s functional relevance to cognitive consequences of perception and motor processes, we address with this study, one such confound, namely the volitional motor report (*35*). Future work aimed at elucidating the causal mechanisms of conscious perception not just in the PFC, but the primate brain in general could greatly benefit from employing direct activation of such perceptually modulated ensembles (*75*). In combination with carefully designed experimental approaches, it could help us both understand the relationship and disentangling the mechanisms underlying consciousness from other cognitive processes (*76*) such as introspection (*27, 77*), attention (*78*), decision making (*79, 80*) or cognitive control (*37, 81*).

## METHODS

### Binocular rivalry paradigm, control paradigms and stimulus presentation

The task consisted of two trial types, namely, the physical alternation (PA) trials and binocular rivalry (BR) trials. Both trial types started with the presentation of a red fixation spot (subtending 0.2 degree of visual angle), cueing the animal to initiate fixation. Upon successful fixation for 300 milliseconds within a fixation window (±8°), a drifting sinusoidal grating (size: 8 degrees (radius), speed: 12-13 degrees/sec, spatial frequency: 0.5 cycles per degree, gratings were drifting vertically up or down) was monocularly presented. During one recording session, we used random dot motion stimulus (field of view 8 degrees (radius), speed 13 degrees/sec, 200 limited lifetime dots and 100% coherence). After one or two seconds, the first stimulus was removed and a second stimulus drifting in the opposite direction was presented in the contralateral eye in PA trials. In BR trials, the second stimulus was added to the contralateral eye without removing the first stimulus. This typically results in perceptual suppression of the first stimulus and is denoted by flash suppression (*20, 42, 43, 48*) in Figure 1A. After this period, visual input alternated physically between oppositely drifting gratings in the PA condition. In the BR condition, the percept of the animal switched endogenously between the discordant visual stimuli, whose temporal histogram could be approximated with a gamma distribution (Figure 1C). The total duration of a single trial/observation period was between 8- 10 seconds. Note that the perception of the animal displayed in Figure 1A is identical in the two conditions, even though the underlying visual input is monocular in PA trials, while it is dichoptic in the case of BR. The eye (where the first stimulus was presented), motion direction (which was presented first) and trial types (PA or BR) were pseudorandomized and balanced in a single dataset. During the entire period of a trial, animals maintained their gaze within a fixation window, which was the same size (±8°) as the stimulus. A liquid reward was given to the animal upon successful maintenance of gaze within the window for the entire trial duration.

The eye movement control experiments using the fixation Off and fixation On paradigms were carried out on a subset of recording sessions (4/6, 2 - H’07, 2 - A’11). Both paradigms consisted of trials, where the macaques were presented with a visual stimulus drifting in one of eight randomly chosen directions for one second (Supplementary Figure 5.1). Each trial started with the presentation of a fixation spot for ∼300 milliseconds, following which a drifting visual stimulus was presented for one second. However, there was one key difference across the two paradigms. During fixation Off, the fixation spot disappeared as soon as the visual stimulus was presented, eliciting OKN and the fixation window (the window within which the animal was required to maintain its gaze) was the entire stimulus (±8°). In contrast, during the fixation On paradigm, a fixation spot overlaid on the stimulus and a smaller fixation window (∼±1 to ±2 degrees) indicated that the monkeys must fixate to complete the trial and receive reward, thus suppressing eye movements. Stimulus parameters were identical to the ones used during the BR paradigm.

Dichoptic visual stimulation was carried out with the aid of a stereoscope and displayed at a resolution of 1280X1024 on the monitors (running at a 60 Hz refresh rate) using a dedicated graphics workstation. Prior to the presentation of the BR paradigm, we carried out a previously described calibration procedure (*48*) which ensured that the stimuli presented on the two monitors through the stereoscope were appropriately aligned and could be fused binocularly. It started with the animal participating in a fixation-saccade task, wherein visual input was at first presented monocularly to the left eye. The task required brief period of fixation on a centrally presented red fixation spot (its location was adjusted according to single eye vergence for each individual monkey), following which a peripheral fixation target was presented in one of eight different directions. Animal was trained to make a saccade to the presented target for obtaining a liquid reward. During this period, the eye position was centered within a fixation window, using a custom designed linear offset amplifier. After this, a second procedure was carried out, wherein the fixation target was first presented to the left eye for a brief duration, after which it was switched off and immediately presented to the right eye. The animal typically responded with a saccade, whose amplitude, provided an estimate of the offset between the fixation spot displayed on the two monitors. This offset was confirmed with several repetitions of this procedure and it served as a correction factor. The visual stimuli were aligned taking into account this correction factor, thus enabling their fusion.

The visual stimuli and the task were designed with an in-house software written in C/Tcl. A QNX real-time operating system (QNX Software Systems) managed the precise temporal presentation of the visual stimuli, and sent digital pulses to the Blackrock recording system. An infrared camera captured eye movements (1kHz sampling rate) with the software iView (SensoriMotoric Instruments GmBH, Germany). Besides monitoring eye movements online, they were also stored for offline analysis in both QNX-based acquisition system as well as Blackrock neural data acquisition system. We used the latter to align the neural data.

### Surgical procedures

Two healthy rhesus monkeys (*Macaca mulatta*), H’07 and A’11 participated in behavioral and electrophysiological recordings. All experiments were approved by the local authorities (Regierungspräsidium, protocol KY6/12 granted to TIP as the principal investigator) and were in full compliance with the guidelines of the European community (EUVD 86/609/EEC) for the care and use of laboratory animals. Each animal was implanted with a cranial headpost (material: titanium) custom designed to fit the skull based upon a high resolution MR scan collected using a 4.7 tesla scanner (Biospec 47/70c; Bruker Medical, Ettlingen, Germany). The headpost implantation was carried out while the animal was under general anesthesia and prior to the beginning of behavioral training in the BR paradigm. Details of the surgical procedures have been previously described (*82*). The MR scan also aided in localizing the inferior convexity of the LPFC. Post behavioral training in the task, the animals underwent another surgery, where a Utah microelectrode array (Blackrock Microsystems, Salt Lake City, Utah USA; (*83*)) was implanted in the PFC. The array had a 10 by 10 electrode configuration and was 4mm by 4 mm in size, with an inter-electrode distance of 400μm and electrode length of 1 mm. We implanted the array ventral to the principal and anterior to the arcuate sulcus, thus aiming to cover a large part of the inferior convexity in the ventrolateral PFC (Figure 2A).

### Electrophysiology data acquisition

All behavioral training and electrophysiological recordings were carried out with the animals seated in a custom designed chair. Data presented here was collected across six recording sessions in two macaques (4 - H’07 and 2 - A’11). Broadband neural signals (0.1 - 30 kHz) were recorded with the Neural Signal Processors (Blackrock Microsystems) and band-pass filtered offline between 0.6 - 3 kHz using a 2nd order Butterworth filter. Spikes were detected with an amplitude threshold set at five times the median absolute deviation (*84*). Any spike events larger than 50 times the mean absolute deviation were discarded. Further, spike events with an inter-spike interval of less than the refractory period of 0.5 ms were also discarded. Events satisfying the aforementioned criterion of threshold and the refractory period were kept for further analysis. Collected spike events were aligned to their minima and 45 samples (1.5 milliseconds) around the peak were extracted for spike sorting. An automatic clustering procedure identified putative single neurons via a Split and Merge Expectation-Maximisation algorithm which fits a mixture of Gaussians on the spike feature data consisting of the first three principal components of the spike waveforms. Inspection and manual cluster cutting was carried out in Klusters (Lynn Hazan, Buzsáki lab, Rutgers, Newark NJ). The details of the spike sorting algorithms have been described elsewhere (*85*). The spiking waveforms, recorded under a given channel, which could not be sorted to a given single unit were collected and denoted as a multi-unit. For the analysis presented in this study, we combined individual single units and multi-units recorded and they are referred to as units.

### Selectivity of unit activity

Each BR trial was visually inspected with the aid of a custom written GUI in MATLAB and the onset and end of a perceptual dominance (during the rivalry phase) was manually marked using the onset of a change in the slow phase of the OKN as a criterion. Two authors VK and AD marked the datasets.

Selectivity of a given unit was assessed separately for PA and BR trials by comparing the spike counts elicited during the presentation (PA) or perception (BR) of downward vs. upward drifting stimuli, using a Wilcoxon rank sum test. For unit selectivity during BR trials, spiking response was aligned to the onset of two events, invoking a perceptual change, namely the (i) onset of flash suppression phase and (ii) onset of a perceptual dominance during spontaneous switches in rivalry. Unit selectivity was similarly assessed during analogous temporal phases of PA trials. The presentation of the second stimulus during PA is temporally corresponding with the presentation of the second stimulus during the BR trial, which constitutes the flash suppression phase. All subsequent stimulus presentations during a PA trial, can be considered equivalent to the perceptual dominance phases during BR. Therefore selectivity of the spiking responses during these periods was computed for assessing unit selectivity during PA trials. Further, among the periods described above, we considered only those epochs during PA and BR trials for computing selectivity, which consisted of perceptual dominance (BR) or monocular presentation (PA) of a given stimulus lasting at least 1000 milliseconds. With respect to perceptual switches, we analyzed transitions, which consisted of at least 1000 milliseconds of clear dominance (judged by a stable OKN pattern), before and after an OKN switch. To compare with PA as close as possible, we analyzed only those transitions with an interval up to 250 milliseconds between the end of the preceding dominance, and the onset of the next. Data was aligned to the onset of the forward dominance. Corresponding temporal phases of stimulus switches during PA trials, included at least one second monocular presentation of a given stimulus followed by the presentation of an oppositely drifting stimulus in the contralateral eye (compared to the preceding visual presentation) for a minimum duration of 1000 milliseconds. Selectivity was assessed both before (-1000 to 0) and after (0 to 1000) the perceptual (BR) and stimulus switches (PA) by collecting all spikes elicited in a 1000 millisecond period. Any relevant figures presented in the main body of the paper were obtained by analyzing PA trials which were aligned to the TTL pulse signaling a stimulus change. In addition, we visually inspected and marked the onset and offset of the visual stimulus during PA trials similarly to the way these episodes were marked for BR trials, based upon the change in the OKN direction. The selectivity analysis (Figure 3) were repeated with PA trials aligned according to this new criterion and the results are presented in Supplementary Figure 3.3, 3.4, 3.5.

### D-prime calculation

For every unit, we computed a preference index denoted as d’, by quantifying the strength of its selectivity during PA and BR trials. It was calculated as follows:

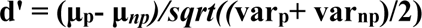

where, **μ_p_** and **μ_np_** is the average spiking response of a given unit during the presentation of its preferred and non-preferred stimulus, calculated over a duration of 1000 milliseconds after a physical stimulus change during PA or a perceptual change during BR trials. The difference between these two quantities is normalized by the square root (**sqrt**) of the average pooled variance (**var_p_** and **var_np_**) of the response distributions.

### Conventional population PSTHs and ensemble PSTHs

Population PSTHs (Figure 3) were computed by averaging the average neural activity of selective units in response to their preferred and non preferred stimuli. The activity of each unit was calculated as the mean response of the unit during specific temporal phases (flash suppression, perceptual dominance and switches) in 50 ms bins during PA or BR trials. For the flash suppression and perceptual dominance phases, we identified all units which displayed significant modulation either during PA or BR trials. With respect to switches, all units which displayed significant modulation (and maintained stimulus preference) before and after a switch during both trial types were identified. In all three cases, the population PSTH was computed by averaging the activity of all units which displayed preference to the same motion direction across PA and BR. In addition, population PSTHs with units significantly selective in the PA or BR conditions were also computed (Supplementary Figures 3.1, 3.4 and 3.2, 3.5, respectively). Population activity related to switches included units which were significantly modulated both before and after the switch for the same motion direction in PA (Supplementary Figure 3.1, 3.4) or BR (Supplementary Figure 3.2, 3.5). Additionally, we generated average ensemble population PSTHs. Here we refer to a population of units displaying preference for the same stimulus as a neuronal ensemble. Population of units which contributed to ensemble PSTHs were identified similarly to conventional population PSTHs. Therefore, the population of units contributing to Figure 3B,C (Switches) and Figure 4A is identical. However, PSTHs were computed differently. First, the activity elicited by all units preferring the downward and upward motion directions were separately averaged for each transition in 50 ms bins, providing a population vector of each neural ensemble for every switch. Next, each of these traces was normalized by subtracting the minimum and dividing it by the maximum activity. Finally, traces were collected across all transitions across datasets and were averaged to generate the average ensemble population PSTHs, presented in Figure 4A. Such an ensemble population PSTH complements the decoding approach, which utilizes the population response on single trials aimed at ascertaining the ongoing sensory input (PA) or perceptual experience (BR). Ensemble population PSTHs for the control paradigms presented in Figure 5A were generated similarly as described above, but without the normalization step. For the ensemble activity related to presentation of the preferred stimulus, all trials where the preferred stimulus of the units comprising the two different ensembles (upward and downward motion) were presented were pooled together and an average was computed. Similarly, all trials where ensemble’s non preferred stimulus was presented were pooled together and averaged for ensemble activity pertaining to the non-preferred stimulus.

### Decoding Analysis

Multivariate pattern analysis was utilized to assess if the spiking activity of neuronal ensembles in the prefrontal cortex contained information about the stimulus on a single transition basis. In this regard, we used a maximum correlation coefficient classifier (*52*) implemented as a part of the neural decoding toolbox (*86*). All the recorded units (N = 987) across the two monkeys were pooled as a pseudopopulation for the decoding analysis pertaining to the BR paradigm (Figure 4). This is similar to previous studies (*52, 87*) where units recorded during independent sessions are treated as simultaneously recorded. Firing rates of each of these units during 15 randomly selected stimulus (PA trials) and perceptual switches (BR trials) were utilized. A z-score normalization (subtracting the mean activity and dividing by the standard deviation) of each unit’s response was done before it participated in the classification procedure in order to assure that units with high spike rates do not influence the decoding procedure disproportionately. We used 15 cross-validation splits, implying that for 14 switches used for training the pattern classifier, one was leaved out to put in the test. This procedure was repeated 50 times (resample runs) to estimate the classification accuracy with a different randomly chosen cross-validation split in each run. All decoding accuracy estimates are zero-one-loss results. Each pixel in cross temporal generalization plots (Figure 4B, 5B and Supplementary Figures 4.1B, 4.2B, 5.4B) depicts the classification accuracy computed with firing rates in 150 ms bins, sampled every 50 ms. This bin duration was chosen, because it has been previously used successfully for decoding visual input from neural activity in the frontal and temporal cortex (*52, 87*). Similar steps as described above were used for decoding analysis in the control paradigms (Figure 5 and Supplementary figure 5.4), with one difference. Units which were significantly modulated in either of the two tasks and preferred the same stimulus (N = 104) participated in the decoding procedure.

Statistical significance of the classification accuracy was estimated using a permutation test, which involved running the decoding analysis on the data with labels shuffled (*51, 86*). This procedure was repeated 500 times with parameters related to binning, cross validation splits as well as resample runs identical to those used for standard decoding of correctly labeled data. The resultant classification accuracies obtained served as a null distribution. If the decoding results obtained without shuffling the labels were greater than all values within the null distribution, they were considered as significant (p<1/500 = 0.002). Significance of decoding accuracy was computed using this procedure for the results presented in figure 4C (also supplementary figure 4.1 and 4.2) and 5C (also supplementary figure 5.4).

### Selection of trials with reduced eye position variance

To create a robust dataset for decoding the visual stimulus during passive observation of monocular stimuli, we only picked those trials corresponding to upward and downward moving gratings where the OKN during passive fixation was relatively flat, i.e. there were no strong drifts in the direction of motion during suppression of eye movements. Firstly, the Y coordinate of the eye movement signal was detrended to remove any DC offset. It was then filtered below 20Hz to remove high-frequency noise and blinks. Next, the double differential was computed and compared to a flat line (i.e. with a slope of 0 and an intercept corresponding to the baseline of the OKN signal) using a least-squared-error minimization method. The sum of the squared error for each trial was collected across all sessions. All those trials whose sum of least squared error was less than the median of the distribution of these errors were selected for further analysis. This method resulted in a selection of trials with reduced variance of the eye movements signal (Supplementary Figure 5.3).

## SUPPLEMENTARY FIGURES

**Supplementary Figure 2.1.**
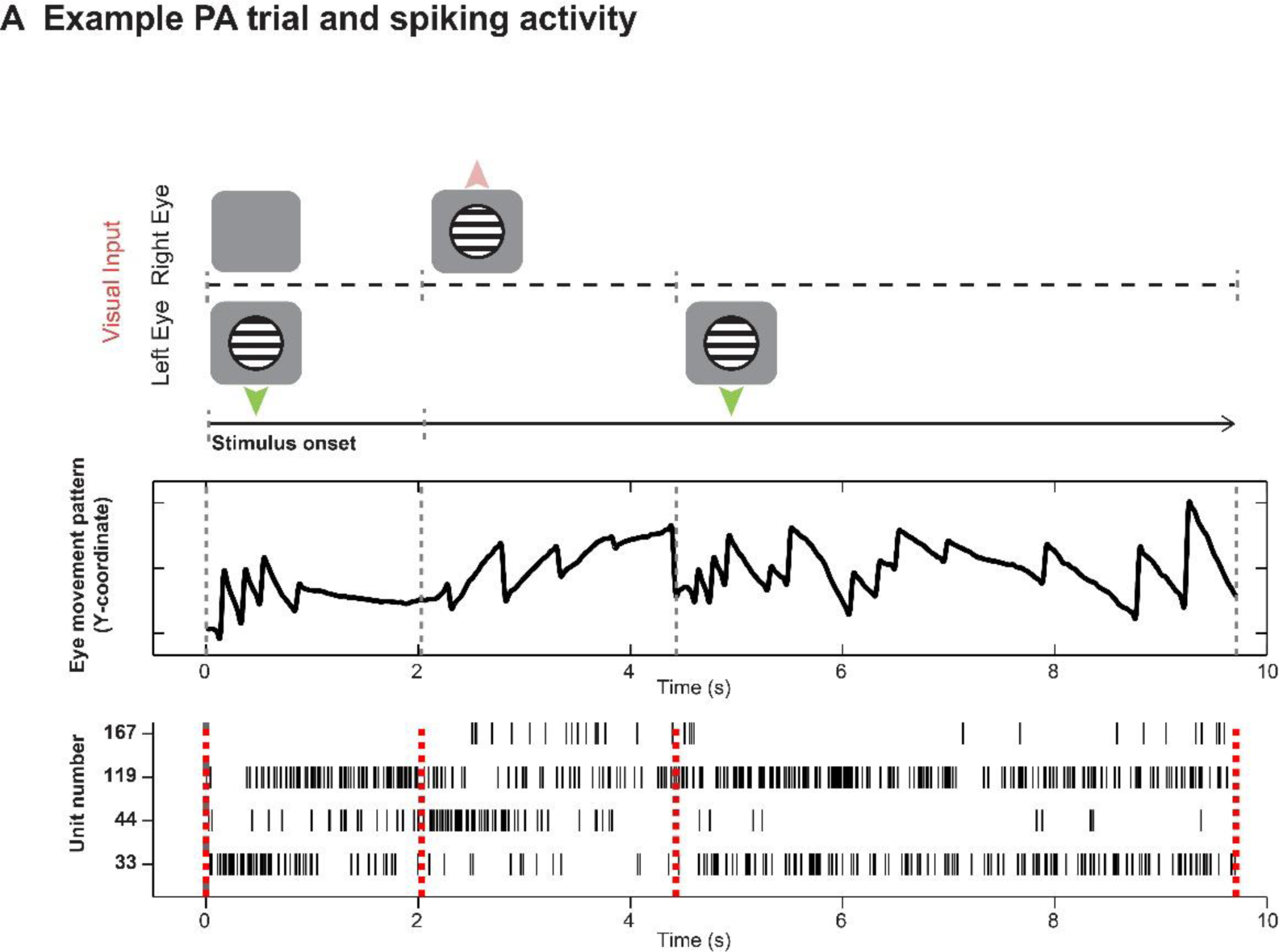
Y-coordinate of the eye movement signal displaying the OKN, and concomitantly recorded spiking activity during an example PA trial for the same units presented in Figure 2B. The trial started with monocular presentation of downward drifting grating. 2000 ms later, the first grating was removed and simultaneously, an upward drifting grating was presented to the contralateral eye. A stimulus switch was externally induced at ∼4500 ms, and a change in the OKN polarity was observed right after. While unit number 33 and 119 display strong spiking activity when downward drifting grating is presented, unit numbers 44 and 167 respond strongly to the presentation of upward drifting grating, thus modulated in a similar way as for perceptual switches during BR in Figure 2B.

**Supplementary Figure 2.2.**
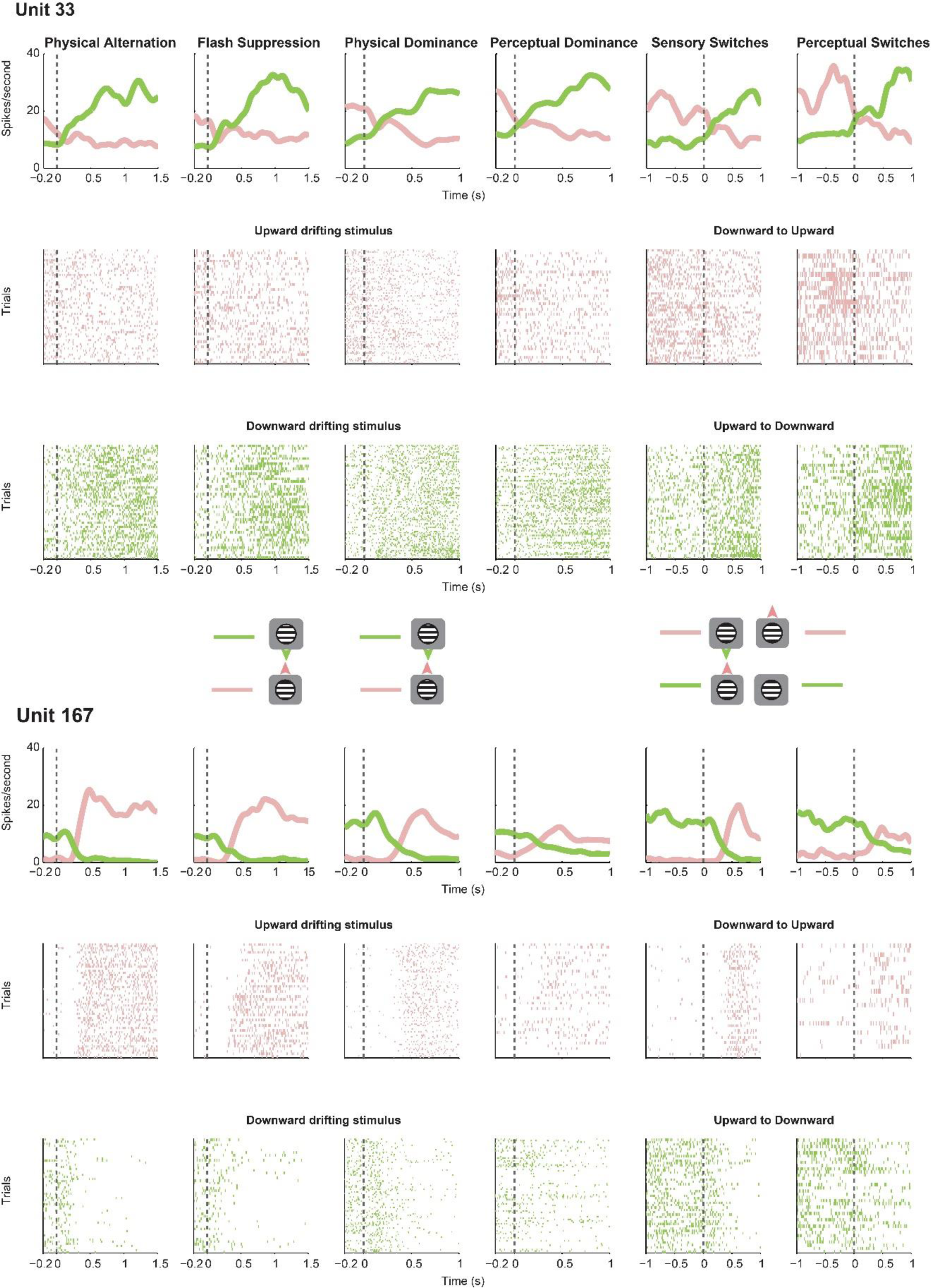
Spike density functions and raster plots, for the units displayed in Figure 2D. Unit 33 displayed stronger activity to grating drifting down, while Unit 167 fired more, when a grating drifting up was presented in PA or perceived in BR. Stronger activity of units is evident also in the spike rasters. With respect to the first four columns of spike rasters: displayed in pink are responses related to grating drifting upwards, while in green is spiking related to the stimulus drifting down. The last two columns display spiking activity as pink rasters for a down to up switch, while in green for an up to down switch.

**Supplementary Figure 2.3.**
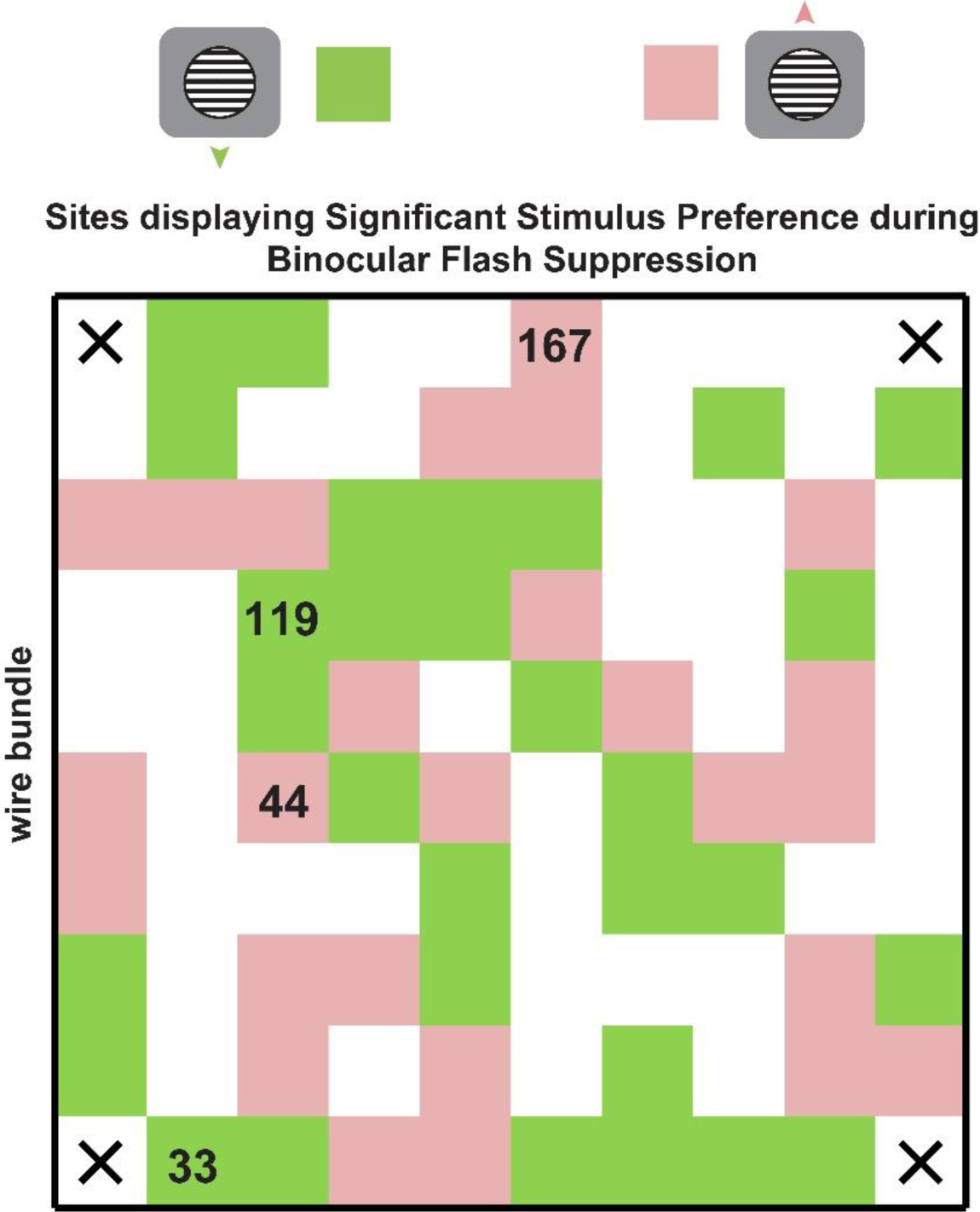
Sites which displayed significant stimulus preference during the flash suppression phase of the BR trials during one recording session is projected back on the array. The numbers denote the location of units displayed in Figure 2B. Green and pink pixels reflect sites, where the spiking activity (unsorted spiking activity recorded from a given electrode), responded more to upward or downward drifting gratings respectively.

**Supplementary Figure 3.1.**
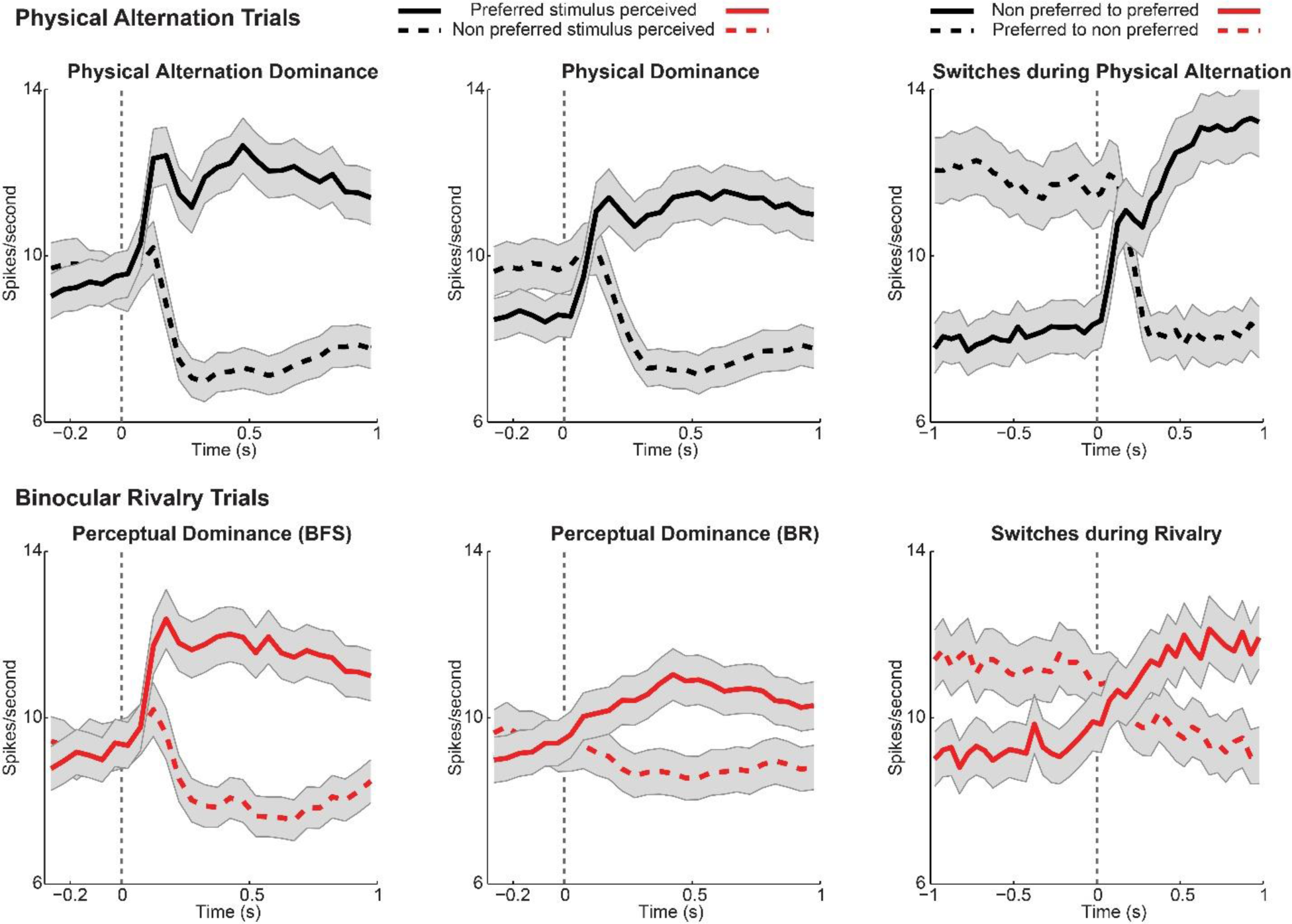
Similar to Figure 3B, the average population spiking activity during PA and BR trials is presented across the various temporal phases of the paradigm (flash suppression, perceptual dominance and switches) for units significantly modulated during PA trials. For switches, selectivity was estimated both before and after the stimulus change. Units significantly modulated more for the same visual stimulus both before and after the stimulus switch were used.

**Supplementary Figure 3.2.**
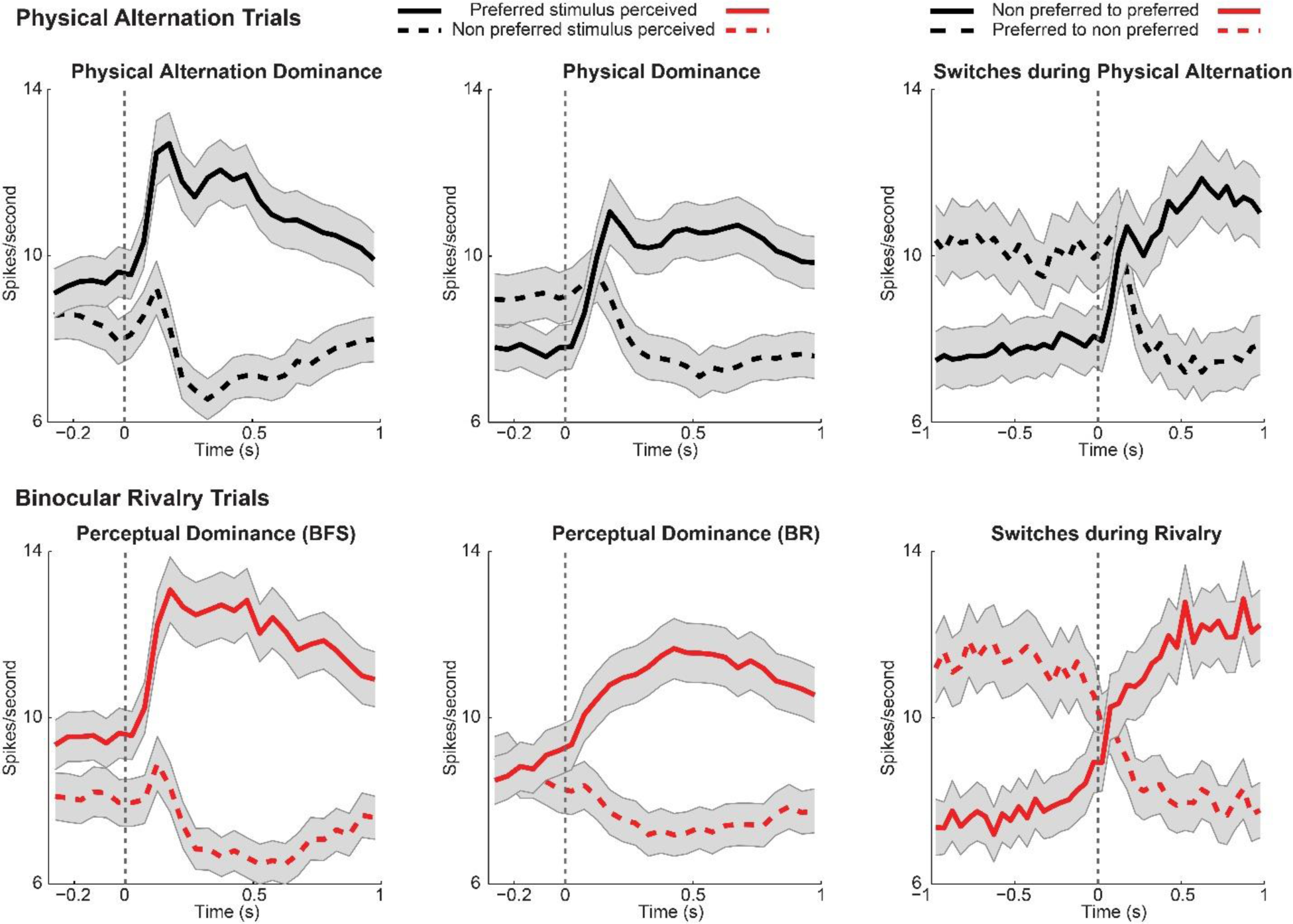
Similar to Figure 3B, presented here is the average population activity during PA and BR trials across the various temporal phases of the paradigm. The population activity was computed using units which were significantly modulated during BR trials. For switches, selectivity was estimated both before and after the perceptual change. Units which were significantly modulated more for the same perceived motion direction both before and after the perceptual transition were used.

**Supplementary Figure 3.3.**
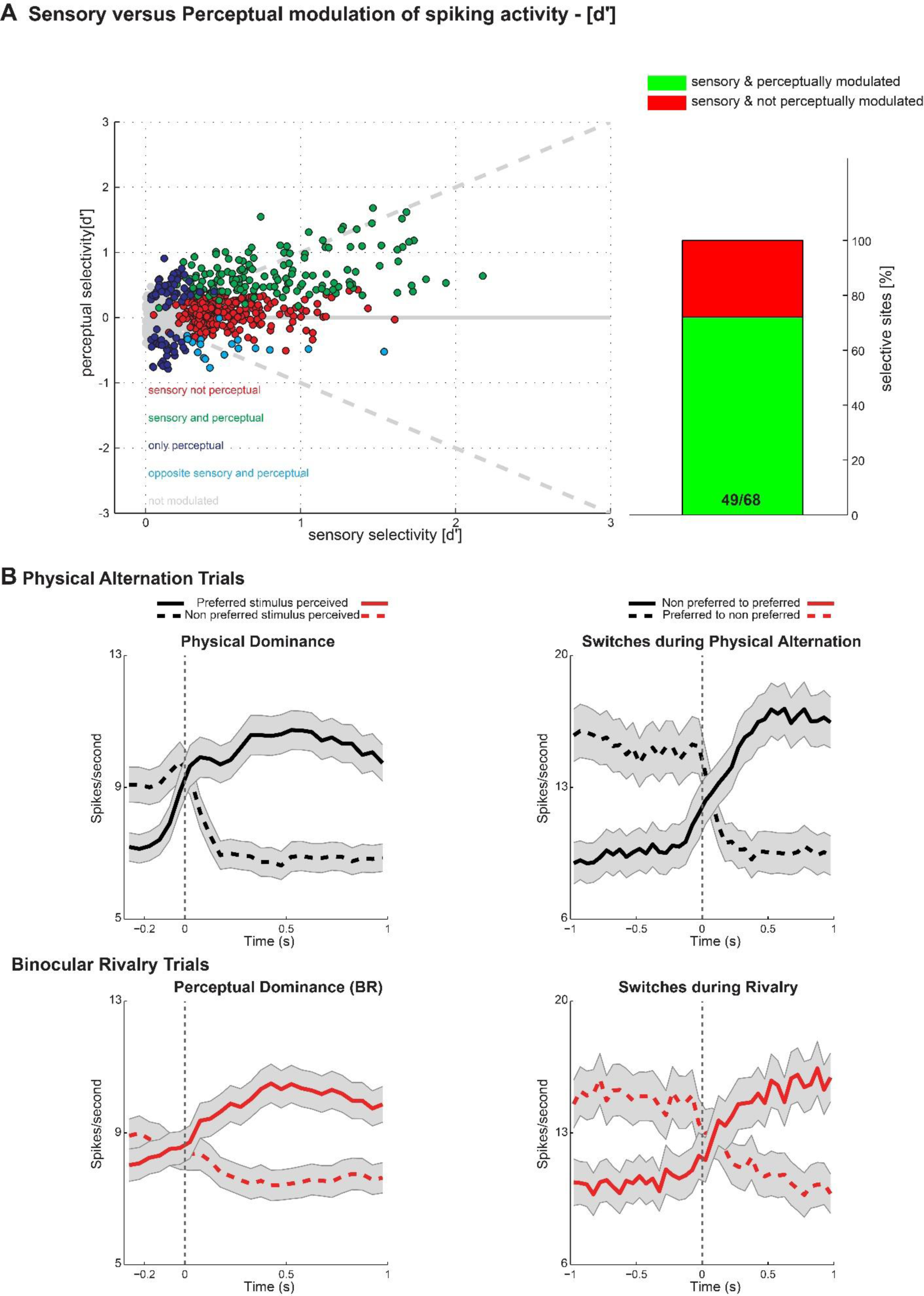
Similar to Figure 3, these plots display the results obtained when the onset and offset of the visual stimulus in physical alternation trials was aligned to the change in OKN (see methods). (A) Scatter plot of sensory versus perceptual preference (d′) for all recorded units is displayed. Each dot denotes a unit. Units showing no significant modulation in PA or BR trials are displayed in grey while those with significant modulation during both conditions are colored green. In red are units which display significant preference only during PA trials. Units displaying significant modulation only during BR trials are displayed in blue, while in cyan are units which fired more when their preferred stimulus was perceptually suppressed. As evident from the scatter plot, the proportion of PA modulated units which are also significantly modulated during BR increase as a function of the strength of sensory selectivity (d′). The right column displays the proportion of PA modulated units with a d′ greater than 1, which were also significantly modulated during perceptual dominance phase in BR trials (green). The right column plots a similar scatter for perceptual dominance phase of the task. (B) Displayed below is the average population spiking activity during PA and BR trials. Similar to Figure 3, population activity was computed by averaging across all units which were significantly modulated during PA or BR trials and preferred the same stimulus. Although the PA trials which participated in this analysis were aligned to change in OKN, the population activity observed across the two trial types was remarkably similar indicating clear and robust perceptual modulation in the units recorded in the vlPFC. Population activity reliably followed phenomenal perception during perceptual transitions brought about both exogenously with flash suppression as well as endogenously driven during BR.

**Supplementary Figure 3.4.**
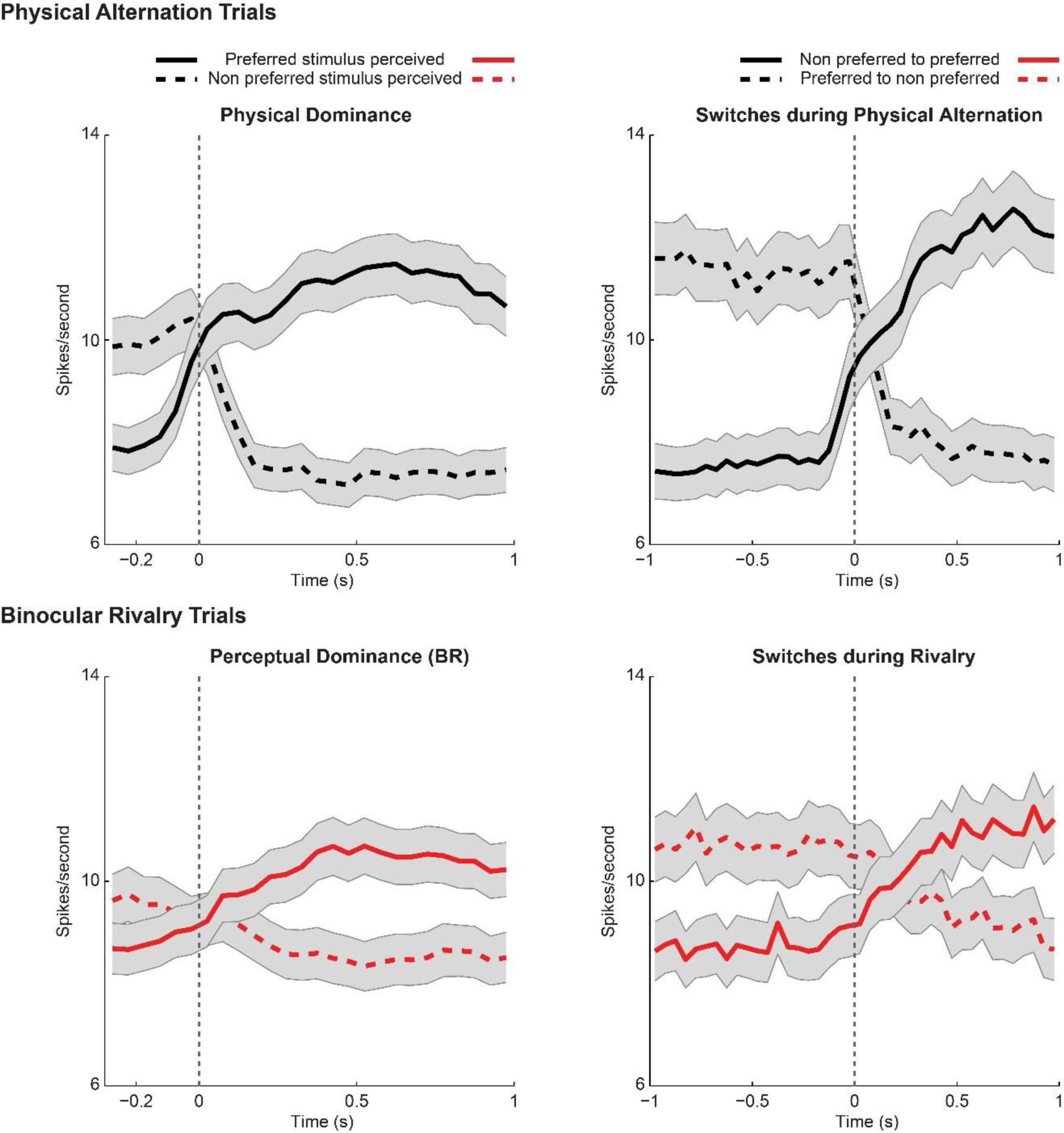
Similar to Figure 3.1 B, results obtained with PA trials aligned to the change in OKN. Presented across three columns is the average population spiking activity during PA and BR trials. The population activity averaged across all units which were significantly modulated during PA trials is plotted here during three temporal phases, namely, the flash suppression phase, the perceptual dominance phase and switches.

**Supplementary Figure 3.5.**
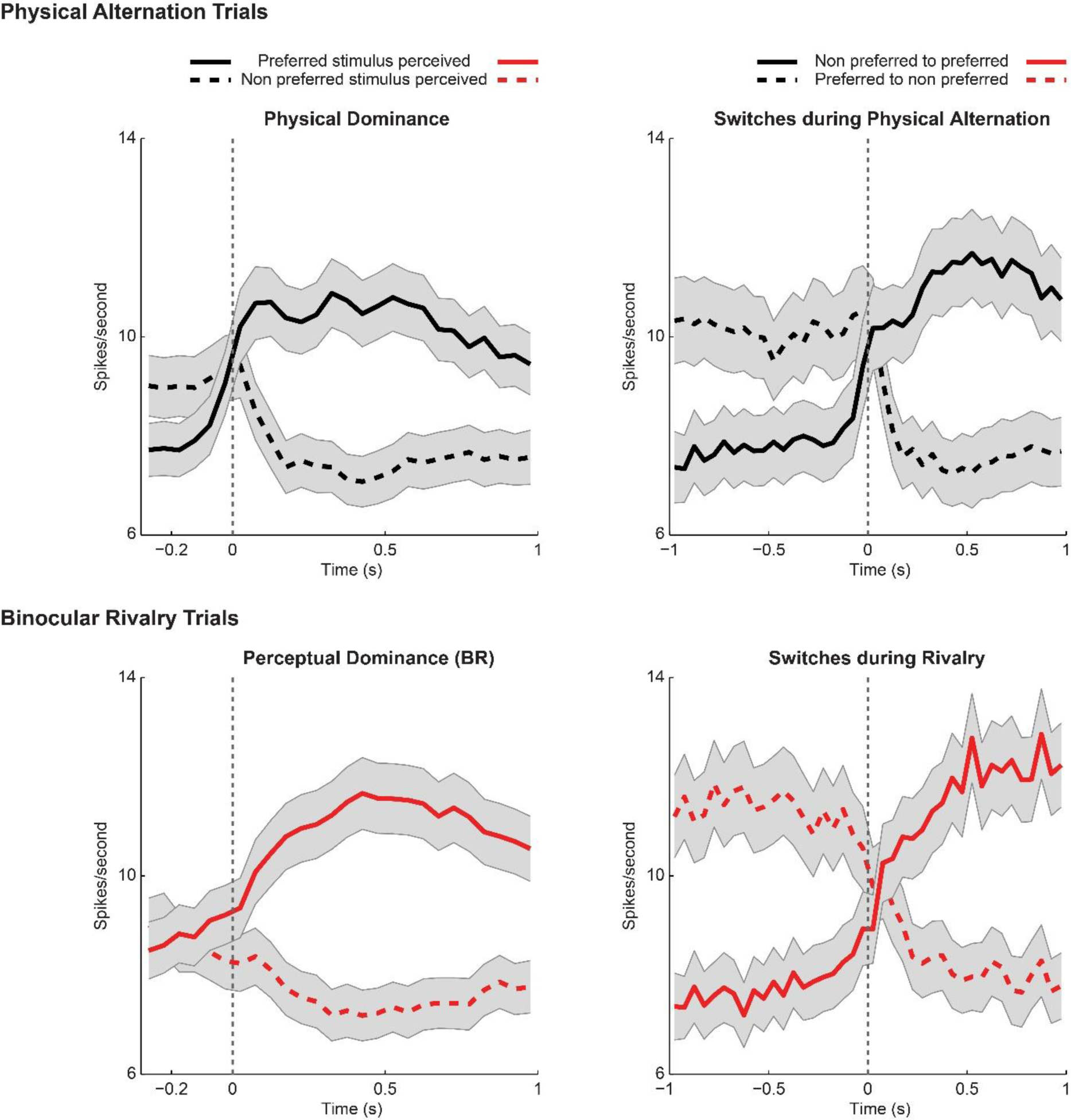
Similar to Figure 3.2, results obtained with manually marked PA trials. Presented across three columns is the average population spiking activity during PA and BR trials. The population activity averaged across all units which were significantly modulated during BR trials is plotted here during three temporal phases, namely, the flash suppression phase, the perceptual dominance phase and switches.

**Supplementary Figure 4.1.**
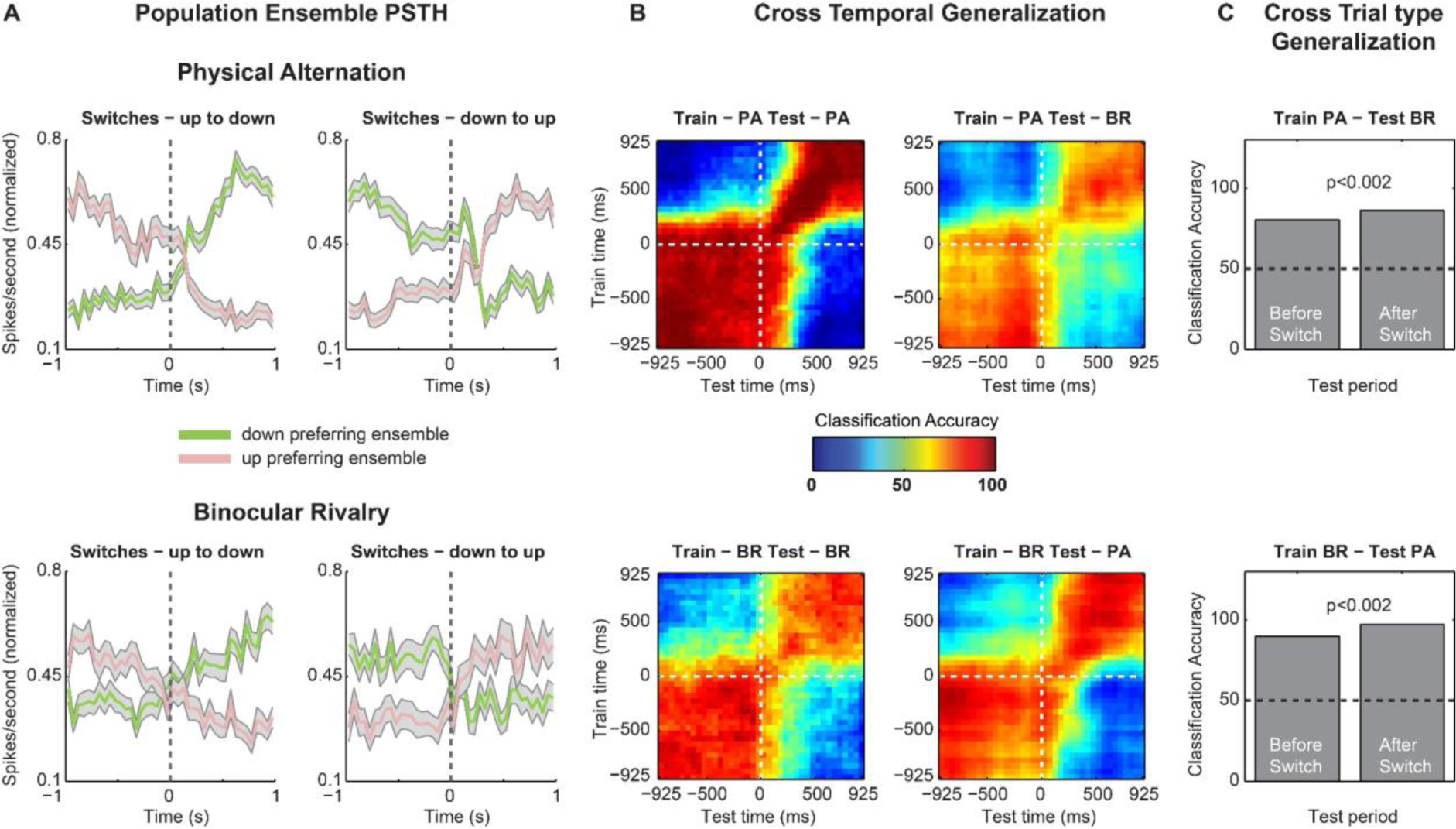
Similar to Figure 4, the results obtained with the multivariate pattern analysis from an individual dataset are displayed. Both stimulus and perceptual contents could be successfully decoded from simultaneously recorded units in an individual dataset.

**Supplementary Figure 4.2.**
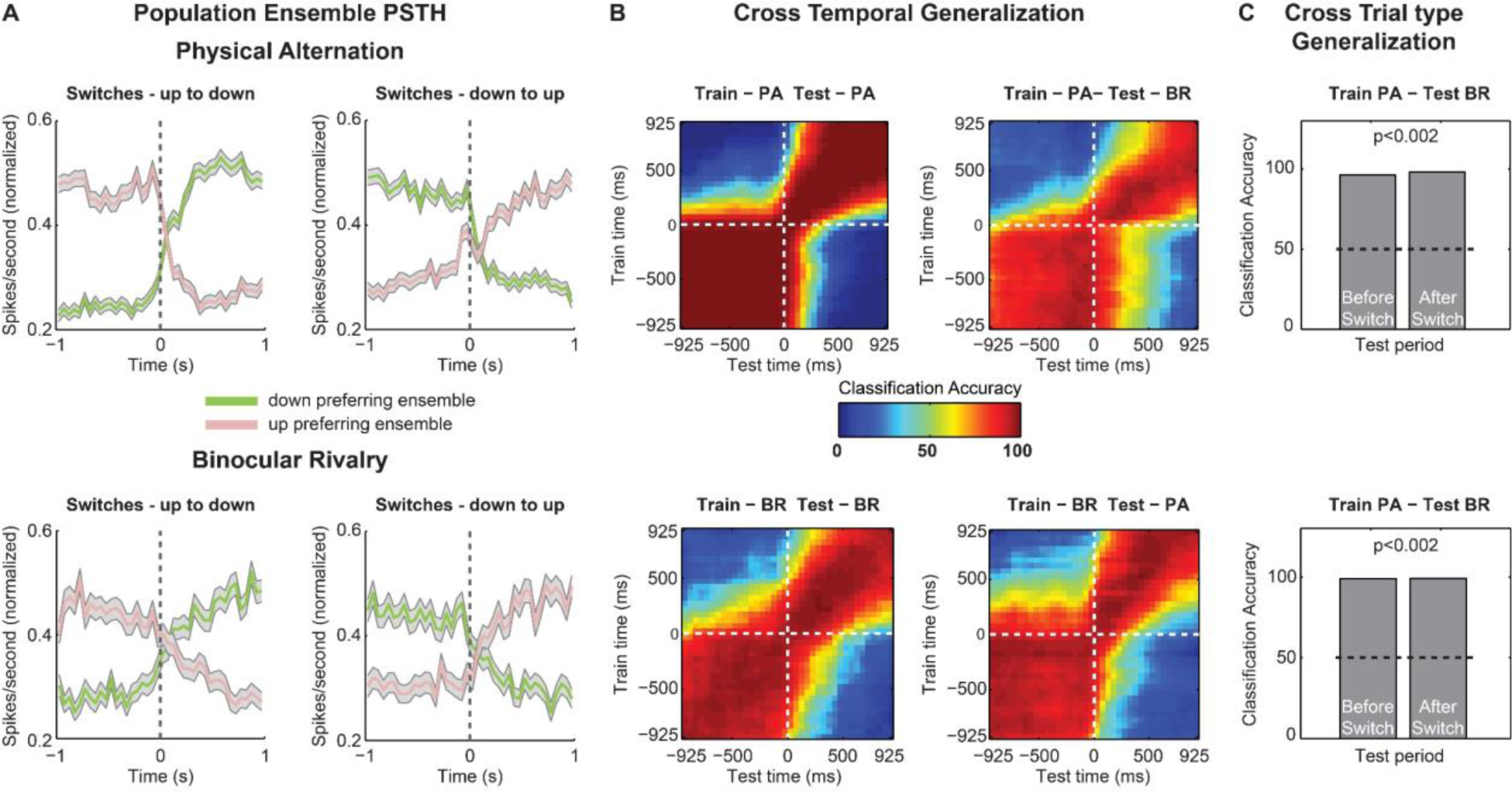
Similar to Figure 4, plotted here are the results obtained with the multivariate pattern analysis, when PA trials were aligned to the change in OKN instead of the TTL pulse (see methods). In is the normalized spiking activity of neuronal ensembles (see methods) during the two different switch types. (B) Cross temporal decoding within and generalization across the two trial types. (C) A cross trial type generalization was carried out over a single temporal window of 400 ms before and after a switch.

**Supplementary Figure 5.1.**
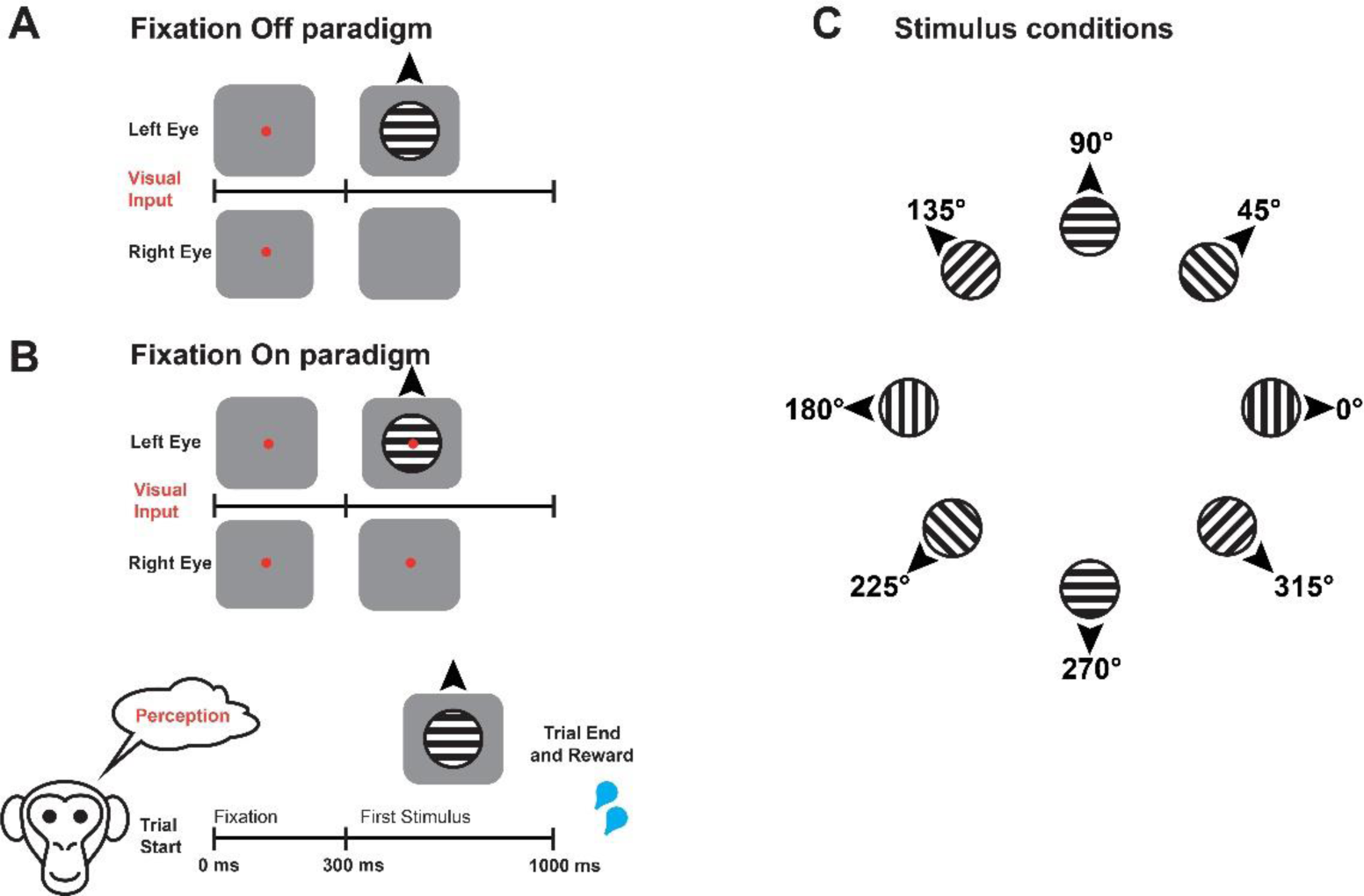
To test the contribution of eye movements on neural activity, animals participated in two control paradigms, namely fixation Off and fixation On, in two different blocks. Both of them started with cueing the animal to fixate for 300 ms, after which a stimulus drifting in a particular direction was presented monocularly. (A) During fixation Off, the fixation spot was removed at the onset of the stimulus, thus inducing optokinetic nystagmus eye movements. (B) During fixation On, the stimulus was presented without removal of the fixation spot, and the animal was required to maintain its gaze within a window (±1 or ±2 degrees) until the end of the trial, in order to receive a juice reward. (C) During both paradigms, on each trial, a stimulus drifting in one of eight different directions was presented.

**Supplementary Figure 5.2.**
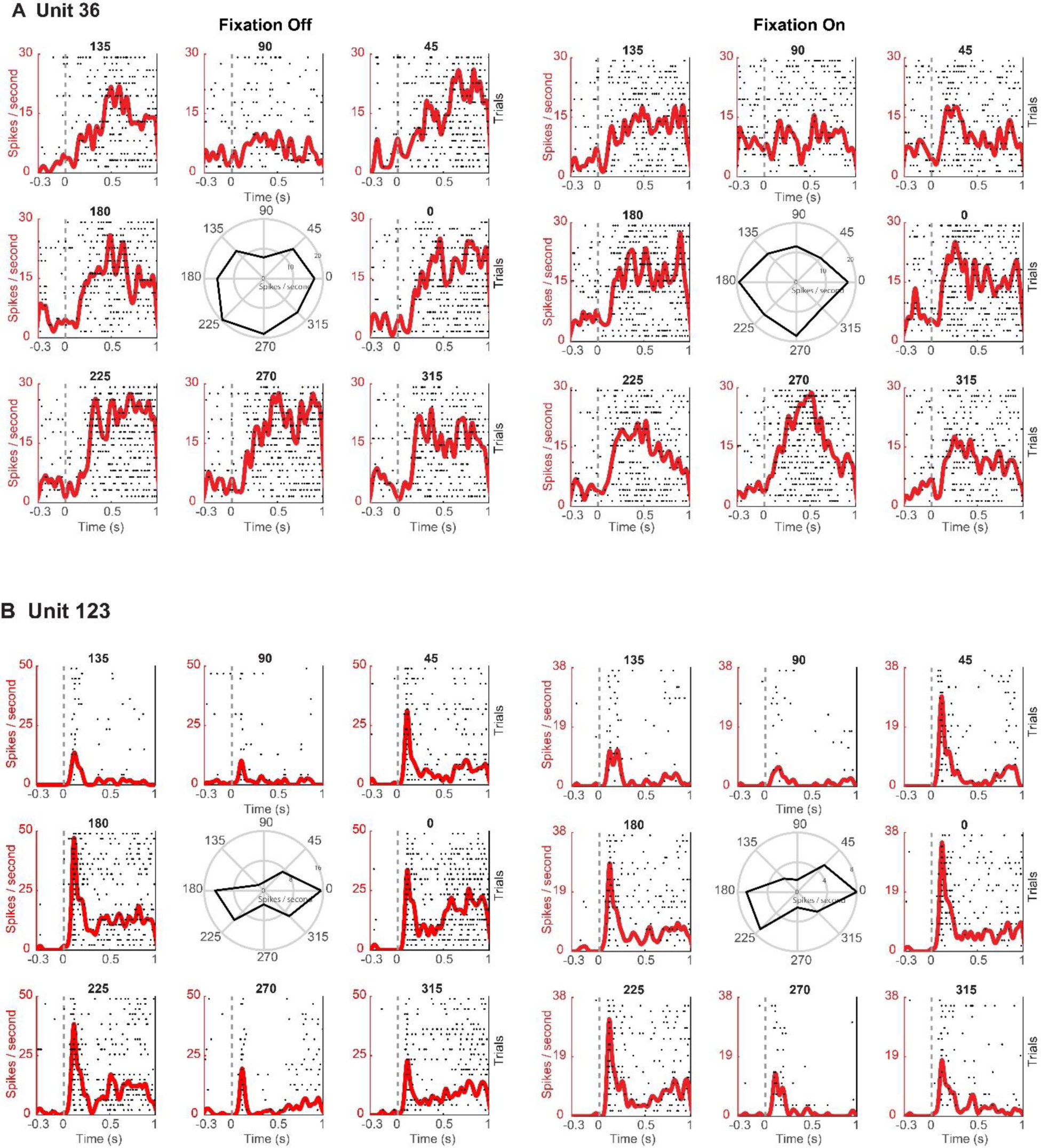
Spike density functions overlaid on spike raster plots for two units during fixation Off and fixation On. Unit activity is presented in response to eight different motion directions. In the middle are polar plots, displaying the tuning curves of each unit (average response of the unit in Hz to drifting gratings in different directions). The presented motion direction was pseudorandomized across trials. Spike rasters are displayed for the first ‘n’ trials presented for every motion direction. Here, n is the minimum number of trials presented to the animal across any motion direction during a given control paradigm. PSTHs and tuning curves were computed taking all trials (of a given motion direction) into account. (A) Unit 36 displays a stronger response to stimulus with motion drifting downwards during both conditions. (B) The unit responds strongly to two opposite directions of motion, thus displaying orientation preference. Note that although the firing rate was higher during the fixation off paradigm, the unit displayed remarkably similar preference in its responses across both paradigms.

**Supplementary Figure 5.3.**
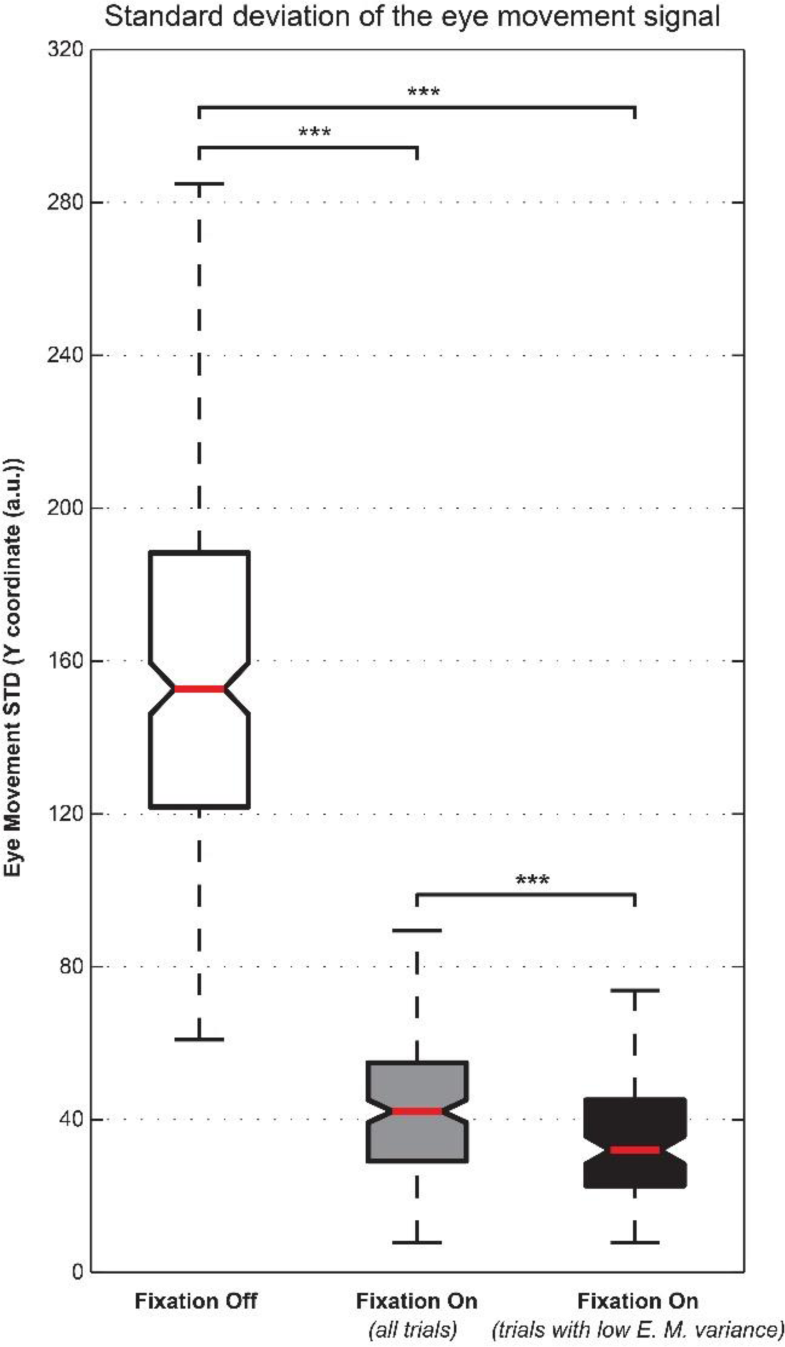
Whisker box plots displaying the distribution of standard deviations (STD) estimated from the y-coordinate of the eye movement signal elicited during individual trials of the two control experiment during fixation Off and fixation On. The STD was computed from the eye movement signal elicited during the time window between 0 (stimulus onset) and 1000 ms (stimulus offset). For fixation On, we used either all trials, or selected trials, which displayed lower variance in the eye movement signal (see methods). The STD of the eye movement signal was significantly reduced (Wilcoxon rank sum test, *** denotes p ≤ 0.001) during fixation On as compared to the fixation Off. The box denotes the 25th (Q1) and 75th percentiles (Q3) of the data, while the red line denotes the median. All adjacent values within Q3 + 1.5×(Q3−Q1) and Q1−1.5×(Q3−Q1) are contained within the upper and lower whisker lengths, respectively. The 95% confidence interval around the median is approximated by the notches, whose edges are calculated as median ± 1.57 (Q3−Q1)/(square root of number of samples).

**Supplementary Figure 5.4.**
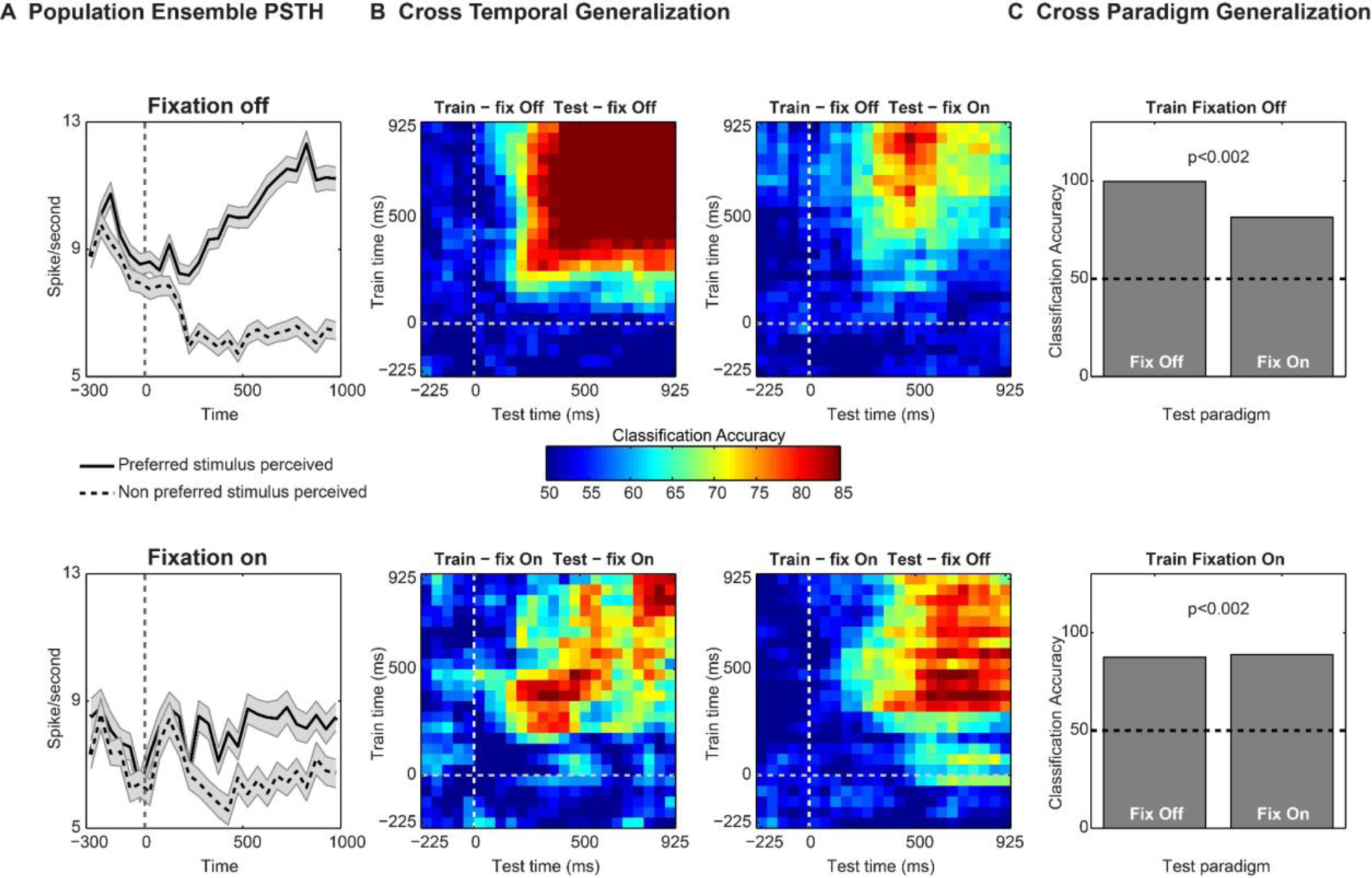
Similar to Figure 5, the figure summarizes the results pertaining to the multivariate pattern analysis assessing the invariance of the population code to motion content during the control experiment. However, only a small selection of the trials from the fixation On paradigm are included, where eye movements were further controlled (see methods and Supplementary Figure 5.3). (A) Population spiking activity (see methods) of prefrontal ensembles is presented during fixation off and fixation On paradigm. The population consisted of units which were significantly modulated in either of the two paradigms, and preferred the same motion direction Cross-temporal decoding of stimulus contents during the two paradigms. Decoding accuracy was tested for each pair of train and test time windows similar to Figure 4 and 5 (C) The cross paradigm invariance of the population code was tested by training a classifier on activity in one paradigm and testing on the other, for a single bin of 400 ms (starting 400 ms post stimulus onset) during the presentation of the visual stimulus. We observed significant (permutation test, p < 0.002) cross-task generalization accuracy, thus suggesting that the underlying code is largely invariant to the presence of large OKN, and encodes stimulus motion contents.

## ACKNOWLEDGMENTS

This study was supported by the Max Planck Society. We would like to thank Prof. Nicho Hatsopoulos and Dr. Yusuke Murayama for help with animal surgery and Axel Oeltermann for his excellent technical help.

## AUTHOR CONTRIBUTIONS

V.K., A.D. and T.I.P. designed the study. V.K., A.D. and S.S. trained animals. V.K. and A.D. performed experiments and collected data, with occasional help from S.S. V.K. and A.D. analyzed the data. S.S. contributed to spike sorting and selectivity analysis of control experiments. M.B. contributed to the decoding analysis. V.K. prepared and arranged the figures in the final format. S.S. provided the MATLAB generated version of the figures displayed in figure 5A, 5.2, 5.3 and 5.4 A. T.I.P. and N.K.L. supervised the study. N.K.L. and J.W. contributed unpublished reagents/analytical tools. N.K.L. provided the support to the group. V.K. and T.I.P. wrote the original manuscript draft. All authors participated in discussion and interpretation of the results and editing the manuscript.

## Bibliography

1. R. Adolphs, The unsolved problems of neuroscience. Trends Cogn Sci (Regul Ed). 19, 173–175 (2015).

2. F. C. Crick, C. Koch, 23 problems in systems neuroscience (Oxford University Press, 2006).

3. G. Miller, What is the biological basis of consciousness? Science. 309, 79 (2005).

4. F. Crick, C. Koch, Towards a neurobiological theory of consciousness. Seminars in the Neurosciences (1990).

5. B. J. Baars, Global workspace theory of consciousness: toward a cognitive neuroscience of human experience. Prog. Brain Res. 150, 45–53 (2005).

6. F. Crick, C. Koch, Consciousness and neuroscience. Cereb. Cortex. 8, 97–107 (1998).

7. D. A. Leopold, N. K. Logothetis, Multistable phenomena: changing views in perception. Trends Cogn Sci (Regul Ed). 3, 254–264 (1999).

8. H. Lau, D. Rosenthal, Empirical support for higher-order theories of conscious awareness. Trends Cogn Sci (Regul Ed). 15, 365–373 (2011).

9. S. Dehaene, J.-P. Changeux, Experimental and theoretical approaches to conscious processing. Neuron. 70, 200–227 (2011).

10. G. Rees, G. Kreiman, C. Koch, Neural correlates of consciousness in humans. Nat. Rev. Neurosci. 3, 261–270 (2002).

11. J. P. Vignal, P. Chauvel, E. Halgren, Localised face processing by the human prefrontal cortex: stimulation-evoked hallucinations of faces. Cogn. Neuropsychol. 17, 281–291 (2000).

12. O. Blanke, T. Landis, M. Seeck, Electrical cortical stimulation of the human prefrontal cortex evokes complex visual hallucinations. Epilepsy Behav. 1, 356–361 (2000).

13. B. Odegaard, R. T. Knight, H. Lau, Should a few null findings falsify prefrontal theories of conscious perception? J. Neurosci. 37, 9593–9602 (2017).

14. A. Del Cul, S. Dehaene, P. Reyes, E. Bravo, A. Slachevsky, Causal role of prefrontal cortex in the threshold for access to consciousness. Brain. 132, 2531–2540 (2009).

15. S. M. Fleming, J. Ryu, J. G. Golfinos, K. E. Blackmon, Domain-specific impairment in metacognitive accuracy following anterior prefrontal lesions. Brain. 137, 2811–2822 (2014).

16. S. M. Szczepanski, R. T. Knight, Insights into human behavior from lesions to the prefrontal cortex. Neuron. 83, 1002–1018 (2014).

17. R. K. Nakamura, M. Mishkin, Chronic “blindness” following lesions of nonvisual cortex in the monkey. Exp. Brain Res. 63, 173–184 (1986).

18. I. Colás et al., Conscious perception in patients with prefrontal damage. Neuropsychologia. 129, 284–293 (2019).

19. B. van Vugt et al., The threshold for conscious report: Signal loss and response bias in visual and frontal cortex. Science. 360, 537–542 (2018).

20. T. I. Panagiotaropoulos, G. Deco, V. Kapoor, N. K. Logothetis, Neuronal discharges and gamma oscillations explicitly reflect visual consciousness in the lateral prefrontal cortex. Neuron. 74, 924–935 (2012).

21. C. Libedinsky, M. Livingstone, Role of prefrontal cortex in conscious visual perception. J. Neurosci. 31, 64–69 (2011).

22. H. Gelbard-Sagiv, L. Mudrik, M. R. Hill, C. Koch, I. Fried, Human single neuron activity precedes emergence of conscious perception. Nat. Commun. 9, 2057 (2018).

23. G. Tononi, M. Boly, M. Massimini, C. Koch, Integrated information theory: from consciousness to its physical substrate. Nat. Rev. Neurosci. 17, 450–461 (2016).

24. C. Koch, What Is Consciousness? Nature. 557, S8–S12 (2018).

25. M. Boly et al., Are the neural correlates of consciousness in the front or in the back of the cerebral cortex? clinical and neuroimaging evidence. J. Neurosci. 37, 9603–9613 (2017).

26. C. Koch, M. Massimini, M. Boly, G. Tononi, Neural correlates of consciousness: progress and problems. Nat. Rev. Neurosci. 17, 307–321 (2016).

27. S. Frässle, J. Sommer, A. Jansen, M. Naber, W. Einhäuser, Binocular rivalry: frontal activity relates to introspection and action but not to perception. J. Neurosci. 34, 1738– 1747 (2014).

28. T. Knapen, J. Brascamp, J. Pearson, R. van Ee, R. Blake, The role of frontal and parietal brain areas in bistable perception. J. Neurosci. 31, 10293–10301 (2011).

29. J. Brascamp, R. Blake, T. Knapen, Negligible fronto-parietal BOLD activity accompanying unreportable switches in bistable perception. Nat. Neurosci. 18, 1672– 1678 (2015).

30. S. Safavi, V. Kapoor, N. K. Logothetis, T. I. Panagiotaropoulos, Is the frontal lobe involved in conscious perception? Front. Psychol. 5, 1063 (2014).

31. N. Zaretskaya, M. Narinyan, Introspection, attention or awareness? The role of the frontal lobe in binocular rivalry. Front. Hum. Neurosci. 8, 527 (2014).

32. J. Brascamp, P. Sterzer, R. Blake, T. Knapen, Multistable perception and the role of the frontoparietal cortex in perceptual inference. Annu. Rev. Psychol. 69, 77–103 (2018).

33. C. Wheatstone, Contributions to the physiology of vision. part the first. on some remarkable, and hitherto unobserved, phenomena of binocular vision. Philosophical Transactions of the Royal Society of London. 128, 371–394 (1838).

34. R. Blake, N. K. Logothetis, Visual competition. Nat. Rev. Neurosci. 3, 13–21 (2002).

35. N. Tsuchiya, M. Wilke, S. Frässle, V. A. F. Lamme, No-Report Paradigms: Extracting the True Neural Correlates of Consciousness. Trends Cogn Sci (Regul Ed). 19, 757–770 (2015).

36. M. A. Pitts, S. Metzler, S. A. Hillyard, Isolating neural correlates of conscious perception from neural correlates of reporting one’s perception. Front. Psychol. 5, 1078 (2014).

37. J. Aru, T. Bachmann, W. Singer, L. Melloni, Distilling the neural correlates of consciousness. Neurosci. Biobehav. Rev. 36, 737–746 (2012).

38. N. K. Logothetis, J. D. Schall, Binocular motion rivalry in macaque monkeys: eye dominance and tracking eye movements. Vision Res. 30, 1409–1419 (1990).

39. M. Wei, F. Sun, The alternation of optokinetic responses driven by moving stimuli in humans. Brain Res. 813, 406–410 (1998).

40. M. Naber, S. Frässle, W. Einhäuser, Perceptual rivalry: reflexes reveal the gradual nature of visual awareness. PLoS ONE. 6, e20910 (2011).

41. R. Fox, S. Todd, L. A. Bettinger, Optokinetic nystagmus as an objective indicator of binocular rivalry. Vision Res. 15, 849–853 (1975).

42. J. M. Wolfe, Reversing ocular dominance and suppression in a single flash. Vision Res. 24, 471–478 (1984).

43. R. W. Lansing, Electroencephalographic correlates of binocular rivalry in man. Science. 146, 1325–1327 (1964).

44. W. J. Levelt, Note on the distribution of dominance times in binocular rivalry. Br. J. Psychol. 58, 143–145 (1967).

45. I. N. Pigarev, G. Rizzolatti, C. Scandolara, Neurons responding to visual stimuli in the frontal lobe of macaque monkeys. Neurosci. Lett. 12, 207–212 (1979).

46. F. A. Wilson, S. P. Scalaidhe, P. S. Goldman-Rakic, Dissociation of object and spatial processing domains in primate prefrontal cortex. Science. 260, 1955–1958 (1993).

47. D. Zaksas, T. Pasternak, Directional signals in the prefrontal cortex and in area MT during a working memory for visual motion task. J. Neurosci. 26, 11726–11742 (2006).

48. G. A. Keliris, N. K. Logothetis, A. S. Tolias, The role of the primary visual cortex in perceptual suppression of salient visual stimuli. J. Neurosci. 30, 12353–12365 (2010).

49. N. K. Logothetis, Single units and conscious vision. Philos. Trans. R. Soc. Lond. B. Biol. Sci. 353, 1801–1818 (1998).

50. D. L. Sheinberg, N. K. Logothetis, The role of temporal cortical areas in perceptual organization. Proc Natl Acad Sci USA. 94, 3408–3413 (1997).

51. E. Meyers, G. Kreiman, in Visual Population Codes, N. Kriegeskorte, G. Kreiman, Eds. (2011), pp. 517–538.

52. E. M. Meyers, D. J. Freedman, G. Kreiman, E. K. Miller, T. Poggio, Dynamic population coding of category information in inferior temporal and prefrontal cortex. J. Neurophysiol. 100, 1407–1419 (2008).

53. M. Dieterich, S. F. Bucher, K. C. Seelos, T. Brandt, Horizontal or vertical optokinetic stimulation activates visual motion-sensitive, ocular motor and vestibular cortex areas with right hemispheric dominance. An fMRI study. Brain. 121 **(Pt** **8**), 1479–1495 (1998).

54. J. N. Kim, M. N. Shadlen, Neural correlates of a decision in the dorsolateral prefrontal cortex of the macaque. Nat. Neurosci. 2, 176–185 (1999).

55. E. Lowet et al., Enhanced Neural Processing by Covert Attention only during Microsaccades Directed toward the Attended Stimulus. Neuron. 99, 207–214.e3 (2018).

56. G. A. Mashour, The controversial correlates of consciousness. Science. 360, 493–494 (2018).

57. T. I. Panagiotaropoulos, V. Kapoor, N. K. Logothetis, Subjective visual perception: from local processing to emergent phenomena of brain activity. Philos. Trans. R. Soc. Lond. B. Biol. Sci. 369, 20130534 (2014).

58. M. C. Schmid, A. Maier, To see or not to see--thalamo-cortical networks during blindsight and perceptual suppression. Prog. Neurobiol. 126, 36–48 (2015).

59. D. A. Leopold, N. K. Logothetis, Activity changes in early visual cortex reflect monkeys’ percepts during binocular rivalry. Nature. 379, 549–553 (1996).

60. N. K. Logothetis, J. D. Schall, Neuronal correlates of subjective visual perception. Science. 245, 761–763 (1989).

61. S. P. O Scalaidhe, F. A. Wilson, P. S. Goldman-Rakic, Areal segregation of face-processing neurons in prefrontal cortex. Science. 278, 1135–1138 (1997).

62. S. P. Scalaidhe, F. A. Wilson, P. S. Goldman-Rakic, Face-selective neurons during passive viewing and working memory performance of rhesus monkeys: evidence for intrinsic specialization of neuronal coding. Cereb. Cortex. 9, 459–475 (1999).

63. M. J. Webster, J. Bachevalier, L. G. Ungerleider, Connections of inferior temporal areas TEO and TE with parietal and frontal cortex in macaque monkeys. Cereb. Cortex. 4, 470–483 (1994).

64. K. G. Thompson, J. D. Schall, The detection of visual signals by macaque frontal eye field during masking. Nat. Neurosci. 2, 283–288 (1999).

65. M. Michel, J. Morales, MINORITY REPORTS: CONSCIOUSNESS AND THE PREFRONTAL CORTEX.

66. J. Morales, H. Lau, The neural correlates of consciousness.

67. M. Rigotti et al., The importance of mixed selectivity in complex cognitive tasks. Nature. 497, 585–590 (2013).

68. V. Mante, D. Sussillo, K. V. Shenoy, W. T. Newsome, Context-dependent computation by recurrent dynamics in prefrontal cortex. Nature. 503, 78–84 (2013).

69. V. Kapoor, M. Besserve, N. K. Logothetis, T. I. Panagiotaropoulos, Parallel and functionally segregated processing of task phase and conscious content in the prefrontal cortex. *Commun*. Biol. 1, 215 (2018).

70. M. Schneider, V. G. Kemper, T. C. Emmerling, F. De Martino, R. Goebel, Columnar clusters in the human motion complex reflect consciously perceived motion axis. Proc Natl Acad Sci USA. 116, 5096–5101 (2019).

71. S. Liu, Q. Yu, P. U. Tse, P. Cavanagh, Neural correlates of the conscious perception of visual location lie outside visual cortex. Curr. Biol. 29, 4036–4044.e4 (2019).

72. R. Brown, H. Lau, J. E. LeDoux, Understanding the Higher-Order Approach to Consciousness. Trends Cogn Sci (Regul Ed). 23, 754–768 (2019).

73. Comparing the major theories of consciousness. - PsycNET, (available at https://psycnet.apa.org/record/2009-19897-077).

74. D. Pal et al., Differential role of prefrontal and parietal cortices in controlling level of consciousness. Curr. Biol. 28, 2145–2152.e5 (2018).

75. J. H. Marshel et al., Cortical layer-specific critical dynamics triggering perception. Science. 365 (2019), doi:10.1126/science.aaw5202.

76. N. Block, What Is Wrong with the No-Report Paradigm and How to Fix It. Trends Cogn Sci (Regul Ed). 23, 1003–1013 (2019).

77. I. I. Goldberg, M. Harel, R. Malach, When the brain loses its self: prefrontal inactivation during sensorimotor processing. Neuron. 50, 329–339 (2006).

78. M. Watanabe et al., Attention but not awareness modulates the BOLD signal in the human V1 during binocular suppression. Science. 334, 829–831 (2011).

79. M. N. Shadlen, R. Kiani, in Characterizing Consciousness: From Cognition to the Clinic?, S. Dehaene, Y. Christen, Eds. (Springer Berlin Heidelberg, Berlin, Heidelberg, 2011), Research and Perspectives in Neurosciences, pp. 27–46.

80. Y. H. R. Kang, F. H. Petzschner, D. M. Wolpert, M. N. Shadlen, Piercing of Consciousness as a Threshold-Crossing Operation. Curr. Biol. 27, 2285–2295.e6 (2017).

81. N. Block, in The Future of the Brain, G. Marcus, J. Freeman, Eds. (Princeton University Press, 2015), Essays by the World’s Leading Neuroscientists, p. 161.

82. N. Logothetis, H. Merkle, M. Augath, T. Trinath, K. Ugurbil, Ultra high-resolution fMRI in monkeys with implanted RF coils. Neuron. 35, 227–242 (2002).

83. E. M. Maynard, C. T. Nordhausen, R. A. Normann, The Utah intracortical Electrode Array: a recording structure for potential brain-computer interfaces. Electroencephalogr. Clin. Neurophysiol. 102, 228–239 (1997).

84. R. Q. Quiroga, Z. Nadasdy, Y. Ben-Shaul, Unsupervised spike detection and sorting with wavelets and superparamagnetic clustering. Neural Comput. 16, 1661–1687 (2004).

85. A. S. Tolias et al., Recording chronically from the same neurons in awake, behaving primates. J. Neurophysiol. 98, 3780–3790 (2007).

86. E. M. Meyers, The neural decoding toolbox. *Front*. Neuroinformatics. 7, 8 (2013).

87. Y. Zhang et al., Object decoding with attention in inferior temporal cortex. Proc Natl Acad Sci USA. 108, 8850–8855 (2011).

